# Molecular basis for the role of Ripr in *Plasmodium falciparum* invasion of human erythrocytes

**DOI:** 10.64898/2026.05.07.723680

**Authors:** Xiao Xiao, Qingmiao Zhou, Danushka S. Marapana, Thomas C. McLean, Rainbow W.B. Chan, Shabih Shakeel, Andrew Leis, Pailene S. Lim, Nicolai C. Jung, Myo T. Naung, Sash Lopaticki, Matthew W. A. Dixon, Alyssa E. Barry, Alan F. Cowman, Stephen W. Scally

## Abstract

*Plasmodium falciparum* causes the majority of severe malaria, and merozoite invasion of erythrocytes is a vulnerable, antibody-accessible step of the blood-stage cycle. PfRipr is an essential component of the PCRCR invasion complex, yet the structural basis for antibody-mediated neutralisation remains unclear. Here, we map inhibitory and non-inhibitory epitopes across PfRipr and show that all potent inhibitors localise to the tail region (EGF6-8). Crystal structures reveal that inhibitory antibodies restrict the flexibility surrounding EGF7. Indeed, EGF7 buried surface area correlates strongly with inhibitory potency, identifying this domain as the principal invasion-inhibitory determinant. Pairwise antibody combinations revealed unexpected synergy, with non-inhibitory mAbs potentiating anti-Rh5 activity. Conditional deletion, sequence replacement or positional swapping of EGF6-8 abolished invasion, demonstrating that both sequence and spatial arrangement are indispensable. These data define EGF7 as a conserved, functionally essential vulnerability and provide a blueprint for rational EGF6-8 immunogen design capable of eliciting *P. falciparum* strain-transcending protection against blood-stage malaria.

## Introduction

Malaria remains a major global health burden: in 2024 an estimated 282 million cases and 610,000 deaths were reported worldwide, with Sub-Saharan Africa bearing the greatest share of disease and mortality. In this region, *Plasmodium falciparum* causes the vast majority of infections and severe illness, accounting for approximately 90% of global malaria deaths, with young children and pregnant women disproportionately affected ^1^. By contrast, *P. vivax* is more geographically widespread across Asia, the Americas and parts of Oceania, sustaining reservoirs of relapsing infection that complicate control efforts ^2^. Together, these epidemiological realities underline the urgent need for improved, durable interventions that reduce morbidity and mortality across diverse transmission settings.

Preventing erythrocyte invasion by the merozoite is a rational blood-stage vaccine strategy because this extracellular phase is directly accessible to antibodies ^3, 4^. The leading blood-stage candidate, PfRh5, has validated this concept: it is highly conserved, functionally essential for erythrocyte invasion ^5, 6^, elicits neutralising antibodies ^7–10^ and has advanced into human trials ^11, 12^. Nevertheless, antigenic diversity and functional redundancy among invasion ligands constrain the breadth and durability of single-antigen approaches, motivating the search for additional conserved, functionally essential targets ^13–15^.

PfRh5 functions as part of PCRCR, a five-membered invasion complex comprising, PfPTRAMP ^16^, PfCSS ^16^, PfRipr ^17^, PfCyRPA ^18, 19^ and PfRh5 ^5, 6^ and whose concerted action is obligatory for PfRh5-mediated engagement of the erythrocyte receptor basigin ^20^ and successful invasion ^16, 21, 22^. While structural and immunological studies of PfRh5 and PfCyRPA have informed antigen design ^16^, PfRipr, a large, multi-EGF-like-domain containing protein that bridges PfCyRPA and the PTRAMP-CSS heterodimer within the complex ^16, 17, 23^, remains comparatively under-characterised. PfRipr is conserved across *Plasmodium* species, elicits invasion-blocking antibodies in immunised rodents and rabbits and in naturally exposed humans, and is implicated in PCRCR-mediated membrane interactions, attributes that make it a compelling vaccine candidate with potential cross-species utility ^20^.

PfRipr plays a central scaffolding role within the PCRCR assembly, bridging PfCyRPA to the PTRAMP-CSS heterodimer and thereby helping to tether the complex to the merozoite surface during invasion ^16, 17, 22–24^. Structurally, PfRipr is divided into a rigid N-terminal core and a flexible C-terminal tail. The core adopts a well-defined fold that engages PfCyRPA and contributes to the extended RCR architecture ^22, 24^, whereas the tail comprises a series of EGF domains (annotated EGF5-10) and a short C-terminal domain (CTD) that has proven conformationally heterogeneous and difficult to visualise in intact complexes ^10, 22, 24^. Antibody mapping and functional assays implicate the tail, particularly its C-terminal EGF modules, as the principal target of invasion-blocking antibodies, indicating that this region is both antibody-accessible and functionally important for PCRCR activity ^10, 25^. Conservation of Ripr orthologues and key structural features across *Plasmodium* species raises the prospect that defined regions of the tail could serve as cross-species vaccine targets ^16, 23^.

Key gaps nonetheless remain, such as which PfRipr antibody epitopes confer merozoite invasion inhibition, how antibody binding alters PCRCR structure or dynamics, and whether discrete PfRipr elements are indispensable for function rather than merely supportive. Addressing these questions is essential to prioritise PfRipr regions for immunogen design and to understand how antibodies can disable the PCRCR assembly during the narrow window of merozoite-erythrocyte interaction. Here, we map neutralising and non-neutralising PfRipr epitopes, determine their structural consequences at atomic resolution, and genetically validate an essential, conserved EGF subdomain, providing a structural and functional blueprint to inform rational PfRipr-based vaccine strategies with the potential to broaden and deepen protection against blood-stage malaria.

## Results

### Growth-inhibitory mAbs target the EGF6-8 subdomains of PfRipr

PfRipr has emerged as an attractive immunogen for blood-stage vaccine development. Immunisation with recombinant PfRipr elicits invasion-inhibitory antibodies that can be more potent than those raised against PfRh5 or PfCyRPA individually ^10^. However, a formulation combining full-length PfRh5, PfCyRPA and PfRipr showed little improvement in antibody-mediated inhibition of merozoite invasion over PfRh5 alone in vaccinated rodents, reinforcing the view that understanding and optimising the inhibitory epitopes of PfRipr was critical to realising its vaccine potential ^26^. Together, these observations motivated a detailed molecular dissection of the PfRipr epitopes that confer antibody-mediated inhibition of *P. falciparum* merozoite invasion.

To delineate the molecular determinants of antibody recognition, a panel of mouse monoclonal antibodies was derived against full-length PfRipr. Twenty-four mAbs were screened by biolayer interferometry (BLI) to select those that bound to distinct epitopes on PfRipr and nine were chosen for further characterisation. These mAbs exhibited broad gene usage for V(D)J recombination (Supplementary Table S1). Additionally, two previously reported mAbs, the inhibitory 1G12 and the non-inhibitory 4A8 ^10^, as well as the cross-species reactive mAb 5B3 raised against the Ripr orthologue from *P. vivax* (PvRipr), previously shown to inhibit *P. knowlesi* (EC_50_ 74 µg/mL) and *P. falciparum* invasion (EC_50_ 2,000 µg/mL), were included in the analysis ^23^.

Nine anti-PfRipr mAbs were assessed individually for their capacity to inhibit *P. falciparum* growth *in vitro* using standard growth inhibition assays (GIAs) (Fig. 1a). At 0.4 mg/mL of mAb, 7B3 demonstrated strong growth inhibition, while 2H1 and 6D2 displayed moderate inhibitory effects. The remaining mAbs 3F5, 9A11, 4H10, 5B11, 7E12 and 8H8 showed no significant inhibitory activity (Fig. 1a). Titration of the inhibitory mAbs 1G12, 7B3, 2H1 and 6D2 demonstrated a dose-dependent inhibition (Fig. 1b). This showed that 1G12 and 7B3 had similar potency of inhibition (EC_50_ 97 and 150 µg/mL respectively), while 2H1 and 6D2 had a significantly lower potency (EC_50_ 1,230 and 2,180 µg/mL).

**Fig. 1.**
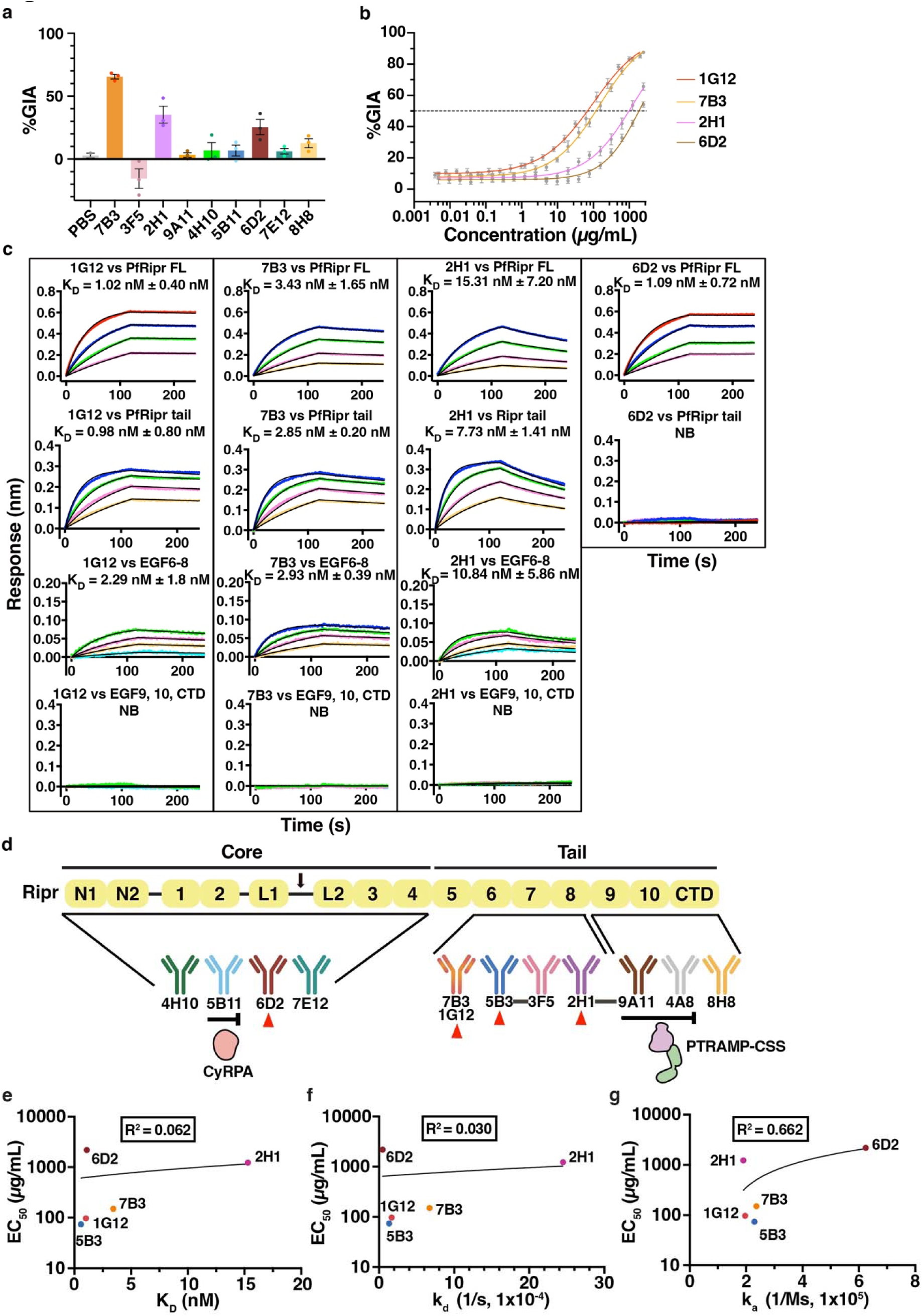
Potent growth-inhibitory anti-PfRipr mAbs target the EGF6-8 subdomains. **a.** *In vitro* GIA using anti-PfRipr mAbs at 0.4 mg/mL. Data are from three independent experiments, except two for PBS vehicle control, each containing three technical replicates. Error bars indicate SEM. **b.** GIA titration with two-fold serial dilution of the inhibitory mAbs 7B3, 2H1 and 6D2 (starting at 2.5 mg/mL) and mAb 1G12 (starting at 2 mg/mL). The %GIA shown is the mean of three independent experiments, each performed in triplicate. Error bars indicate SEM. Half Maximal Effective Concentration (EC_50_) was determined using non-linear regression four-parameter analysis via Prismv9 (GraphPad). All GIAs were performed with 3D7 *P. falciparum* parasites. **c.** Kinetics studies and binding affinities of inhibitory anti-PfRipr mAbs to full-length PfRipr (FL), PfRipr tail (PfRipr^tail^), EGF6-8 (PfRipr^EGF6-8^) and EGF9,10,CTD (PfRipr^EGF9-10-CTD^) measured using BLI. Concentrations of antigen are colored as follows: 500 nM (orange), 250 nM (red), 125 nM (dark blue), 62.5 nM (green), 31.25 nM (pink), 15.63 nM (yellow) and 7.81 nM (light blue). Representative sensorgrams (association and dissociation) and 1:1 model best fit (black) was representative of two independent measurements. Reported equilibrium dissociation constant (K_D_) indicates mean **±** SEM. NB: No Binding. **d.** Schematic of PfRipr domain architecture showing core and tail regions containing two N-terminal domains (N1 and N2), two lectin-like domains (L1 and L2), EGF-like domains numbered 1-10 and the C-terminal domain (CTD). Black arrow represents Plasmepsin X cleavage site. Anti-PfRipr mAb epitope bins are shown below, grouped according to kinetics and competition assays. Bins connected by black lines indicate competition. The mAbs that block complex formation are indicated with the blocking antigen written below (5B11: PfCyRPA; 9A11 and 4A8: PfPTRAMP-CSS). mAbs that exhibit growth-inhibitory activity are indicated with a red triangle. **e.** K_D_ from kinetics studies against PfRipr FL plotted against EC_50_ (log scale) for inhibitory anti-PfRipr mAbs (R² = 0.062). **f.** Dissociation rate constant (k_d_) against PfRipr FL plotted against EC_50_ (log scale) (R² = 0.030). **g.** Association rate constant (k_a_) against PfRipr FL plotted against EC_50_ (log scale) (R² = 0.662). **e-g.** Cross-reactive mAb 5B3 was also included ^23^, with kinetics parameters against PkRipr protein reported in Extended Data Fig. 5a. EC_50_ for 5B3 shown was determined from two-cycle GIA titration against *P. knowlesi* ^23^. All other mAbs shown were against *P. falciparum* in one-cycle GIA. Correlation determined using Shapiro-Wilk test of normality (⍰=0.05) and simple linear regression via Prism v10 (GraphPad; Source Data File).

The position of the binding epitopes for inhibitory and non-inhibitory mAbs were mapped across the full length PfRipr (FL) protein using discrete recombinant subdomains that included the tail (PfRipr^tail^), EGF6-8 (PfRipr^EGF6-8^), EGF9-10-CTD (PfRipr^EGF9-10-CTD^), CTD (PfRipr^CTD^), EGF6 (PfRipr^EGF6^), EGF6-7 (PfRipr^EGF6-7^), EGF7-8 (PfRipr^EGF7-8^), and EGF8 (PfRipr^EGF8^) (Extended Data Fig. 1). This positioned the inhibitory epitopes for mAbs 1G12, 7B3 and 2H1 within the PfRipr tail, specifically the EGF6-8 region (Fig. 1c, d and Extended Data Fig. 3), while 3F5 also engaged EGF6-8 (Fig. 1d, Extended Data Fig. 2 and 3). The less potent mAb 6D2 bound PfRipr FL but not the isolated tail, consistent with an epitope in the core domain (Fig. 1c and d). Similarly, the non-inhibitory mAbs 4H10, 5B11 and 7E12 bound to the core whilst 9A11 bound EGF9-10, and 4A8/8H8 the CTD (Fig. 1d, Extended Data Fig. 2 and 3).

Competition experiments using BLI with the mAbs resolved ten discrete epitope bins on full-length PfRipr and clarified relationships between these assignments (Fig. 1d and Extended Data Fig. 4). The most potent inhibitory mAbs 1G12 and 7B3 occupy the same bin consistent with overlapping epitopes. Similarly, 5B3 competes with 3F5 suggesting partial overlap of each binding epitope. The mAbs 2H1 and 9A11 showed competitive binding to PfRipr despite domain mapping placing the 2H1 epitope within the EGF6-8 region and the 9A11 epitope within the EGF9-10 region (Fig. 1d and Extended Data Fig. 4). This indicates that the epitopes of these mAbs are closely apposed in the context of the flexible tail in three-dimensional space, and binding of one mAb could interfere with accessibility for the other.

The competition assays identified mAbs that functionally block binding of PfRipr to its binding partners PfCyRPA or PfPTRAMP-CSS in the PCRCR complex (Fig. 1d and Extended Data Fig. 4) ^16^. Specifically, mAb 5B11 inhibited PfCyRPA binding to PfRipr, while 9A11 and 4A8 prevented association of PfPTRAMP-CSS with PfRipr (Extended Data Fig. 4). The binning and domain-mapping analyses indicated that these non-inhibitory antibodies target residues at, or immediately adjacent to, the relevant protein-protein interfaces, consistent with previous work ^16^ and with a model in which the PCRCR complex is pre-assembled in merozoite micronemes prior to surface exposure ^27^. Under such a model, antibodies that block formation of individual binary contacts may have limited impact on invasion because those interactions are already established prior to antibody binding. Accordingly, the blockade of PfCyRPA or PfPTRAMP-CSS association by mAbs did not reliably predict growth inhibition. These data therefore emphasise that precise epitope position and the mechanism of antibody engagement, not simply disruption of single binary interactions, determine neutralising activity and should guide the selection of antibody combinations and immunogen design.

To determine whether epitope location or binding kinetics better explain functional differences, antibody-antigen kinetics were quantified by BLI. All mAbs bound with high affinity, with equilibrium dissociation constants (K_D_) spanning 0.33 nM (5B11) to 36.05 nM (9A11) (Extended Data Fig. 5a-c). Correlation analyses with EC_50_ values showed no meaningful relationship between K_D_ (R² = 0.062), dissociation rate constant k_d_ (R² = 0.030) or association rate constant k_a_ (R² = 0.663) and inhibitory potency (Fig. 1e-g). Crucially, several very high-affinity mAbs (for example 5B11, K_D_ 0.33 nM; 4A8, K_D_ 0.60 nM) were non-inhibitory (Extended Data Fig. 5a), indicating that affinity *per se* does not predict activity.

### Anti-PfRipr and anti-PfRh5 antibody combinations reveal synergistic inhibition

Having established that potent neutralising antibodies converge on EGF7, we next asked whether simultaneously targeting distinct epitopes across PfRipr, and across the broader PCRCR complex, could enhance inhibition of *P. falciparum* merozoite invasion. Pairwise combinatorial growth inhibition assays were performed using a panel comprising EGF6-8-directed mAbs (7B3, 2H1, 3F5, 5B3), the moderately inhibitory core-domain binder 6D2 and two anti-PfRh5 antibodies including the potent neutraliser R5.016 and the non-neutralising potentiator R5.011 ^28^. R5.011 has previously been shown to potentiate inhibitory anti-Rh5 antibodies by slowing erythrocyte entry and thereby extending the window for inhibitory antibodies to act ^28^. Synergy was assessed using the Bliss independence model of additivity ^25, 28, 29^.

The interaction landscape spanned antagonism, additivity and strong synergy (Fig. 2a). Strikingly, the non-inhibitory EGF7-8 binder 3F5, which produced negligible %GIA alone, showed powerful potentiation when combined with anti-Rh5 antibodies R5.016 and R5.011 (Fig. 2a-c and Extended Data Fig. 6). Similarly, the weakly inhibitory core-domain binder 6D2 synergised robustly with 3F5 and with both anti-Rh5 mAbs (Fig. 2a-c and Extended Data Fig. 6). For example, titration of 6D2 in the presence of a constant 500 µg/mL of R5.011 improved the 6D2 EC_50_ from 2,180 µg/mL to ∼6.1 µg/mL; 6D2 alone produced only 4% GIA at 125 µg/mL, yet this increased to 75% when combined with 500 µg/mL R5.011 and to 60% when combined with 4 µg/mL R5.016 (Fig. 2b and Extended Data Fig. 6). These results mirror the R5.011 paradigm and demonstrate that antibodies with little or no standalone inhibitory activity can become potent synergistic partners when combined with antibodies targeting distinct sites on the PCRCR complex.

**Fig. 2.**
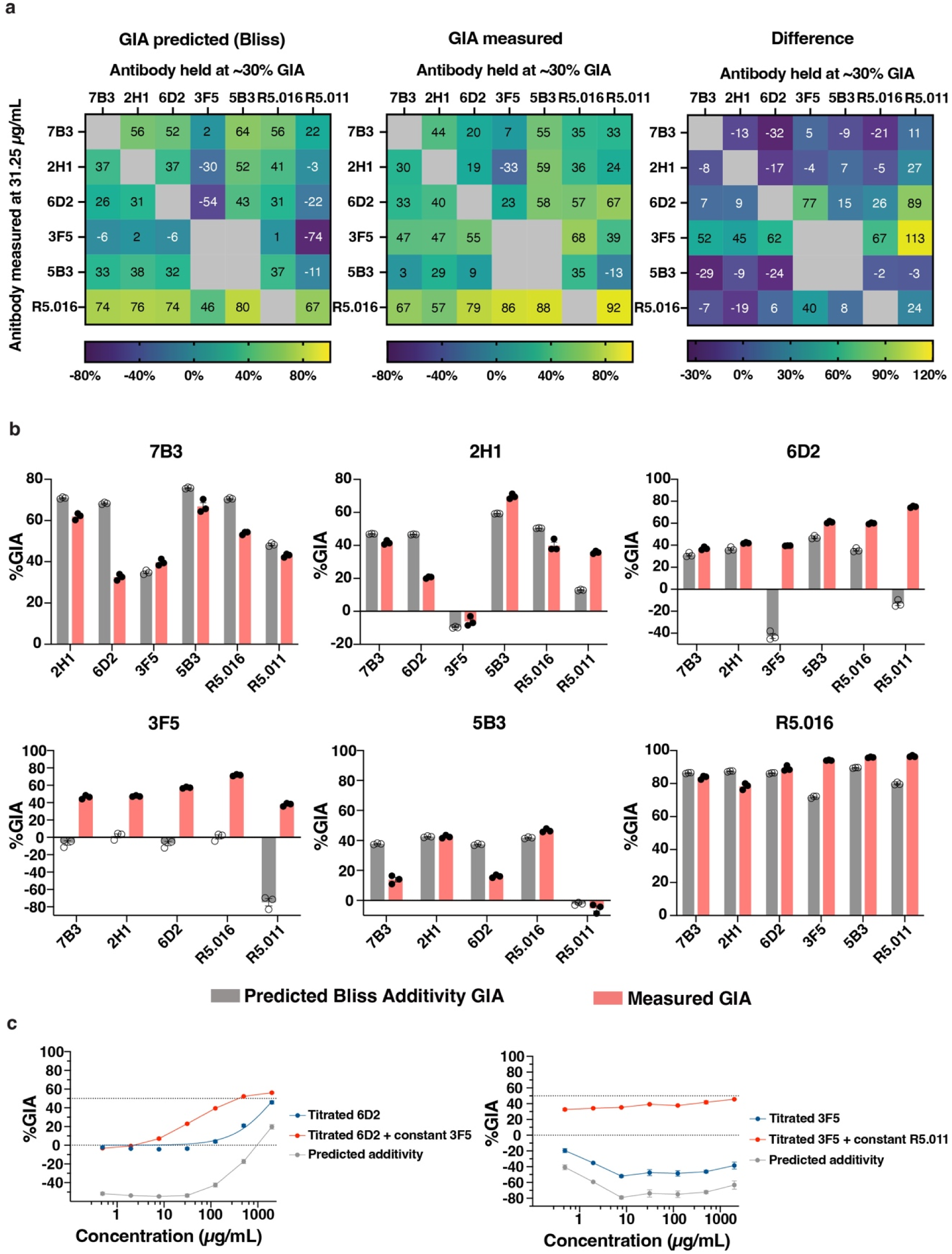
Anti-PfRipr mAbs display a broad synergy profile. **a.** Heatmap of pairwise combination GIA analysis for anti-PfRipr mAbs (7B3, 2H1, 6D2, 3F5, 5B3) and anti-PfRh5 mAbs (R5.016, R5.011) measured at 31.25 µg/mL per mAb, with partner mAb concentration held constant achieving approximately 30% GIA, except for 3F5 and R5.011 which were held at 500 µg/mL due to 3F5 and R5.011 alone having no GIA activity. R5.011 was performed in one-way combinations only. Left: predicted %GIA based on Bliss additivity of each combination; centre: measured %GIA of each combination; right: %GIA difference (measured minus predicted). Combinations with ≥15% GIA above predicted were classified as synergistic; ≥15% below predicted as antagonistic and otherwise as additive. Values represent the mean of a triplicate measurement. **b.** Predicted Bliss additivity GIA (grey) compared with measured GIA (red) for pairwise mAb combinations. One mAb was held at a concentration achieving approximately 30% GIA (x-axis) while the partner mAb was held at 125 µg/mL. 3F5 and R5.011 were held at 500 µg/mL as they have no GIA activity alone. Technical replicates are plotted with error bars indicating SEM. **c.** Exemplar combinatorial GIA titration curves showing synergy. Left: 6D2 titrated in four-fold seven-step serial dilution from 2 mg/mL with constant 3F5 held at 500 µg/mL; right: 3F5 titrated in four-fold seven-step serial dilution from 2 mg/mL with constant R5.011 held at 500 µg/mL. Both pairs show measured %GIA well above predicted additivity across the concentration range, confirming robust synergy. Individual points are the mean of a triplicate measurement with error bars indicating SEM.

In contrast, the inhibitory mAbs (7B3, 2H1, 5B3) were generally additive and in some pairings mildly antagonistic when combined with other antibodies held constant (Fig. 2a,b and Extended Data Fig. 6). The anti-PfRh5 mAb R5.011 tended to be broadly synergistic when held constant, except in combinations with 7B3 and 5B3, which were largely additive. Synergy with R5.016 depended on the partner mAb and could be synergistic, additive or antagonistic (Fig. 2a, b and Extended Data Fig. 6). Collectively, these data show that the anti-PfRipr mAb panel displays a broad and varied synergy profile, both within the panel and in combination with anti-PfRh5 antibodies, indicating that the inhibitory outcome of antibody combinations targeting the PCRCR complex cannot be predicted from the activity of individual antibodies alone and that the relationship between epitope location, antibody class and combinatorial effect is complex.

### Inhibitory anti-Ripr antibodies target EGF7

To define the molecular basis of inhibition by anti-Ripr antibodies, the crystal structures of Fabs from three inhibitory mAbs (1G12, 2H1, 7B3) and one non-inhibitory mAb (3F5) in complex with PfRipr EGF subdomains were determined. Specifically, structures were obtained for 1G12 and 7B3 in complex with PfRipr^EGF6-8^, 2H1 in complex with PfRipr^EGF6-7^, and 3F5 in complex with PfRipr^EGF7-8^ (PDB IDs: 24HU, 24HV, 24HW, 24HY respectively) (Fig. 3a). Owing to the inherent flexibility of the PfRipr tail, not all regions of the EGF domains were resolved in every structure. Nevertheless, Fab binding stabilised discrete regions of PfRipr, enabling detailed structural interpretation of epitope recognition and insight into their potential inhibitory mechanisms.

**Fig. 3.**
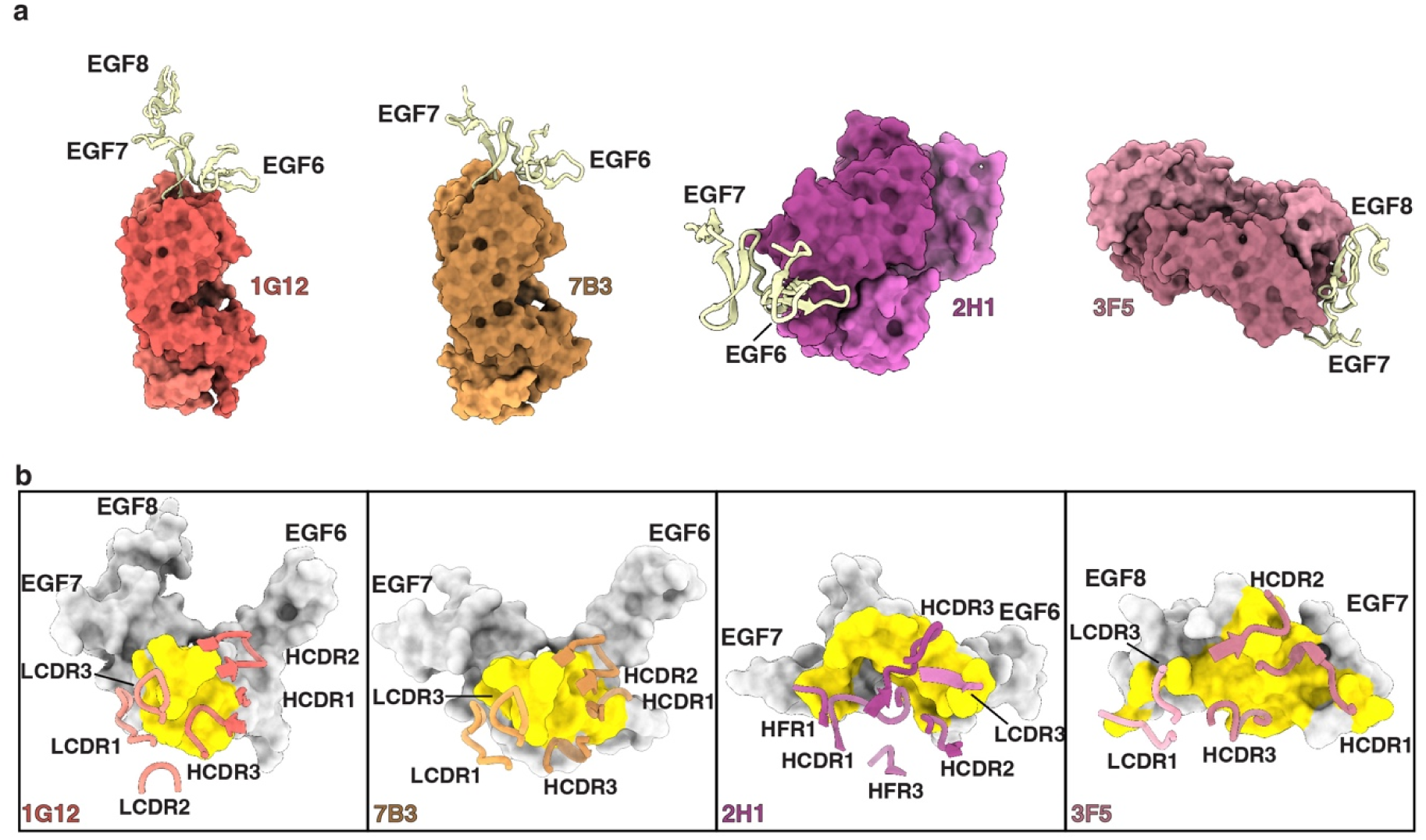
Inhibitory anti-PfRipr antibodies bind PfRipr^EGF7^. **a.** From left to right: crystal structures of PfRipr subdomains (yellow) in complex with Fab fragments of 1G12 (EGF6-8), 7B3 (EGF6-7), 2H1 (EGF6-7) and 3F5 (EGF7-8). Heavy chains of Fab fragments are coloured in darker shade while light chains are coloured in lighter shade (PDB IDs: 24HU, 24HV, 24HW, 24HY respectively). **b.** Corresponding structures from (a) with antibody CDR contacts mapped onto the surface of EGF subdomains (grey). Epitopes are coloured yellow. CDRs are coloured according to (a).

The two most potent anti-PfRipr mAbs, 1G12 and 7B3, recognised a largely overlapping epitope centred on the EGF7 domain, consistent with the epitope binning data (Fig. 3a and b). Notably, while 1G12 and 7B3 share a highly similar light chain differing by only 7 residues, their heavy chains are substantially divergent, differing by 37 residues (Supplementary Table S1). Despite this, their complementarity-determining regions (CDRs) adopted largely similar topology overall, contributing almost equally to their total buried surface area (BSA) of 679 Å^2^ and 680 Å^2^, respectively (Supplementary Table S8). Furthermore, while both mAbs primarily interact with the EGF7 domain, particularly the loop between the main β-strands and a short 3_10_-helix at the beginning of the domain, both also contact the linker region between EGF6-7, with 7B3 also interacting with EGF6 via its CDR1 and CDR2 (Fig. 3a, b, Supplementary Table S3 and S4). Together, these structures provide an example of two divergent mAbs engaging a highly overlapping inhibitory epitope in a conserved manner.

Although 2H1 also bound to EGF6 and EGF7, it recognised a distinct epitope from those of 1G12 and 7B3, consistent with epitope binning data (Fig. 3a and b). In particular, 2H1 primarily engaged the second β-hairpin of EGF6, the EGF6 linker, and the 3_10_-helix of EGF7, almost exclusively via interactions mediated by its heavy chain (Fig. 3a, b, Supplementary Table S5 and S8). In contrast, the non-inhibitory mAb 3F5 bound a distinct epitope, centred on one face of EGF8, as well as the second β-hairpin of EGF7 and the linker region between EGF7-8 (Fig. 3a, b and Supplementary Table S6). Notably, 3F5 had the largest BSA of the four structures, largely mediated by its CDRs, at 820 Å^2^ (Fig. 3b and Supplementary Table S8).

To evaluate antigenic conservation, 20,646 *P. falciparum* field isolate genomes were analysed to map observed sequence variation on the PfRipr structures (Extended Data Fig. 7a). Sequence analysis of the PfRipr^EGF6-8^ domain revealed a high degree of sequence conservation, with complete conservation across the four antibody binding sites (Extended Data Fig. 7b). This exceptional conservation across circulating parasite strains makes EGF6-8 an attractive target for rational immunogen design aimed at eliciting broadly neutralising, *P. falciparum* strain-transcending antibodies that inhibit erythrocyte invasion.

### Inhibitory mechanism of anti-Ripr mAbs

To identify structural features that distinguish inhibitory from non-inhibitory anti-Ripr mAbs, the four Fab-PfRipr structures, together with the crystal structure of 5B3-PvRipr^EGF7-8^ (PDB ID: 9YIO) ^23^ were superimposed using shared EGF subdomains (Fig. 4a). In this configuration, 5B3 sterically clashes with 3F5 on EGF7-8, consistent with epitope binning data (Fig. 4a). Each mAb family adopted a distinct angle of approach to the superimposed complex (Fig. 4b). Notably, despite occupying distinct non-competitive epitopes, 2H1 and 1G12/7B3 induced an identical conformational kink at the junction between EGF6 and EGF7. Indeed, relative to the AlphaFold3-predicted model of the PfRipr tail, the inhibitory anti-PfRipr mAbs (1G12, 7B3 and 2H1) induced a pronounced kink corresponding to a 72° angle between EGF6 and EGF7, compared with the 121° angle predicted for the unbound, elongated PfRipr tail ^30^ (Fig. 4c). In contrast, the non-inhibitory mAb 3F5 bound an extended epitope spanning EGF7 and EGF8, with the angle between these domains approximating 103°, in close agreement with the AlphaFold3 prediction (Fig. 4c) ^30^. By comparison, 5B3, a cross-reactive mAb that potently inhibits *P. knowlesi* and weakly inhibits *P. falciparum*, did not kink the EGF7-8 junction, instead inducing a more open conformation of 125°, 22° greater than that observed in the 3F5-bound conformation (Fig. 4c). Together, these observations suggests that inhibitory antibodies distort the architecture of the Ripr tail and restrict EGF flexibility in a manner likely to disrupt PCRCR complex function.

**Fig. 4.**
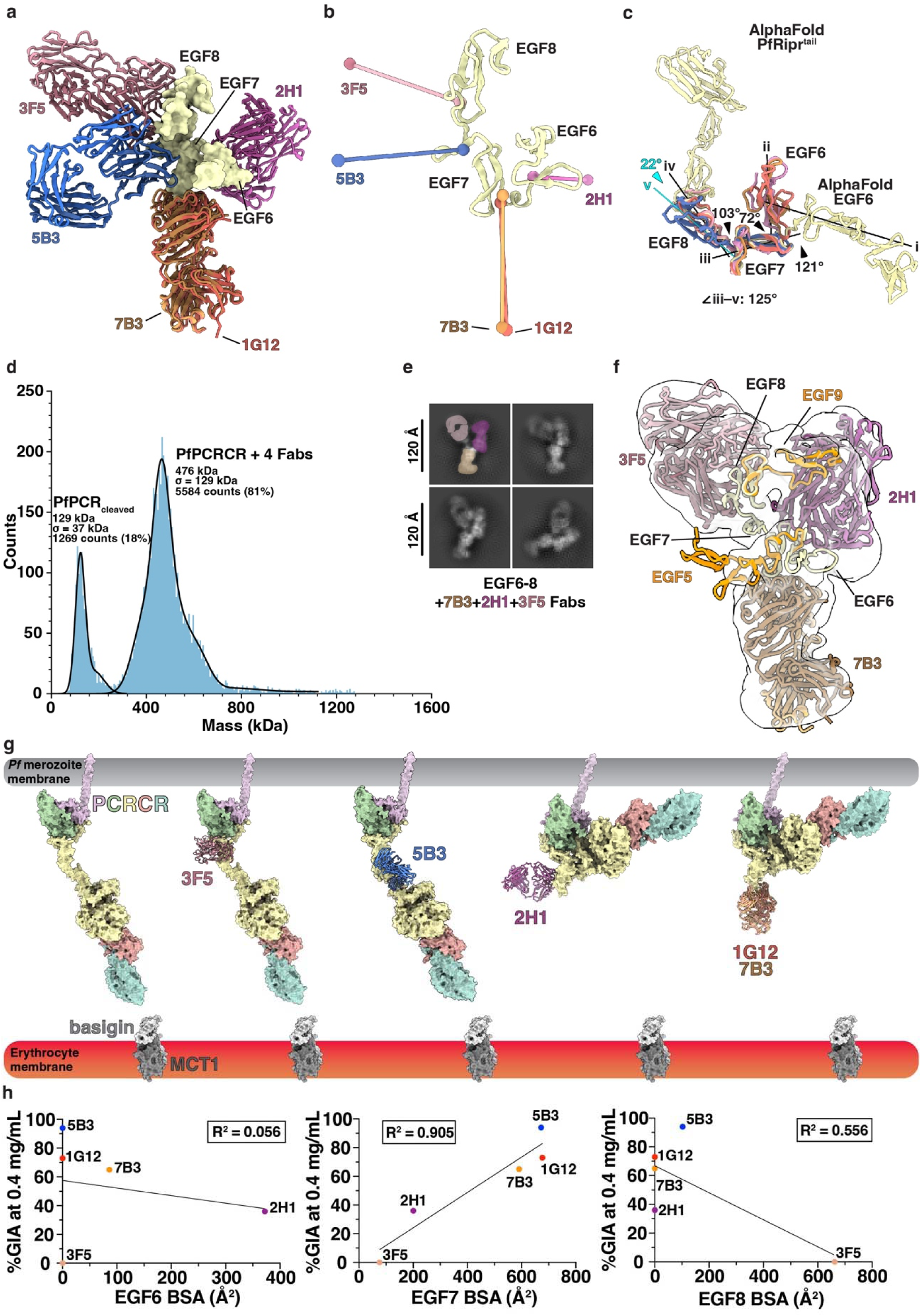
Inhibitory anti-PfRipr antibodies distort the PfRipr tail. **a.** Structural superposition of all four anti-PfRipr Fab onto PfRipr^EGF6-8^ (yellow), with the addition of anti-PvRipr^EGF7-8^ 5B3 Fab (blue; PDB ID: 9YIO) ^23^. Crystal structures were aligned by the sequences of common EGF subdomains in the complexes using PyMOL and ChimeraX. **b.** Comparison of the angle of approach. The centre of mass for both the Fab fragments and interacting EGF residues are shown as spheres and coloured as in (a). Centre of mass determined in ChimeraX. **c.** Overlay of EGF subdomains (coloured according to the Fab fragments in (a)) from each crystal structure onto the AlphaFold3-predicted PfRipr^tail^ (yellow), aligned by the EGF7 subdomain. 1G12, 7B3 and 2H1 induce a 72° kink between EGF6 and EGF7, compared with a 121° angle in the AlphaFold3 prediction of the unbound PfRipr^tail^. 3F5 does not induce angular changes; the EGF7–EGF8 angle remains at approximately 103°, agreeing with AlphaFold3 prediction. 5B3 induces a 125° obtuse extension between EGF7 and EGF8. i-v represent the axes of EGF subdomains in the following order: EGF6 (AlphaFold3), EGF6 (1G12, 7B3 and 2H1), EGF7 (all structures), EGF8 (AlphaFold3, 3F5 and 1G12) and EGF8 (5B3). Axes definition and angles between axes determined in ChimeraX. **d.** Mass distribution of stabilised PfPCRCR-Fabs co-complex after pre-incubation and SEC as measured by mass photometry. Histogram data are shown in blue and Gaussian curve fit in black. **e.** Cryo-EM 2D class averages of the 7B3+2H1+3F5 Fabs-PfRipr complex, coloured according to the superposition in (a). **f.** Cryo-EM volume (transparent surface, EMDB-80770) containing a composite model (cartoon) of 7B3, 2H1 and 3F5 Fabs bound to PfRipr^EGF6-8^ (PDB IDs: 24HV, 24HW, 24HY), and the AlphaFold3 predictions of EGF5 and EGF9 subdomains (orange). **g.** From left to right: crystal structures of 3F5, 5B3, 2H1, 1G12 and 7B3 Fabs (cartoon) aligned with a composite model of the PfPCRCR complex (surface), comprising the cryo-EM structure of PfRh5-CyRPA-Ripr^core^ (PDB ID: 8CDD) and the AlphaFold3-predicted PfPTRAMP-CSS-Ripr^tail^. The receptor basigin and MCT1 (PDB ID: 7CKR) are located on the erythrocyte membrane. **h.** Correlation between buried surface area (BSA, Å^2^) of individual EGF6-8 subdomains contacted by anti-PfRipr^EGF6-8^ mAbs and their respective %GIA at 0.4 mg/mL. Left: EGF6 BSA (R² = 0.056); centre: EGF7 BSA (R² = 0.905); right: EGF8 BSA (R² = 0.556). BSA values for 5B3 are from the previously determined 5B3-PvRipr^EGF7-8^ crystal structure ^23^. %GIA at 0.4 mg/mL for 5B3 was determined from two-cycle GIA titration against *P. knowlesi* ^23^. Correlation was determined using Shapiro-Wilk test of normality (⍰=0.05) and simple linear regression via Prism v10 (GraphPad; Source Data File). BSA was determined by PISA^40^.

To understand how anti-Ripr mAbs engage the PfRipr tail in the broader context of the PCRCR complex, cryo-EM single particle analysis was performed on the assembled complex. To overcome the low affinity of PfPTRAMP-CSS for PfRipr in *P. falciparum*, a stabilised co-complex was engineered by introducing a disulfide-linkage between PfPTRAMP and PfRipr. These proteins were co-expressed with PfCSS and subsequently co-complexed with PfCyRPA and PfRh5, together with the anti-PfRipr Fabs 7B3, 2H1, 3F5 and 6D2 (Fig. 4d and Extended Data Fig. 8a). Although the full complex could not be resolved, we observed densities corresponding to 3 out of 4 Fabs in complex with PfRipr EGF6-8 subdomains. This map at 3.5 Å resolution suffered heavily from orientation bias as evident from cFAR value of 0.02 and particle distribution plot (Extended Data Fig. 8b). Therefore, to reduce the effect of orientation bias, we sub-selected particles using ‘rebalance orientation’ job in cryoSparc and further performed non-uniform refinement, which resulted in a 5.14 Å resolution map with cFAR value of 0.10 (EMDB-80770). This cryo-EM map enabled rigid-body fitting of the crystal structures of 7B3, 2H1 and 3F5 Fab-PfRipr, supporting the validity of the structural superposition and the simultaneous engagement of PfRipr EGF6-8 subdomains by multiple antibodies within the PCRCR complex (Fig. 4e, f). The AlphaFold3 models of EGF5 and EGF9 subdomains were further fitted into the cryo-EM density, revealing that EGF9 wraps around EGF6-8 to form a U-shaped fold (Fig. 4f). The relative positions of EGF5-9 subdomains, not previously visualised, provide new insight into the overall architecture and folded conformation of the PfRipr tail in the context of multiple Fab engagements. With the crystal structure superposition of the assembled co-complex now validated by the cryo-EM model, we propose a model for the mechanism of antibody-mediated inhibition (Fig. 4g). In its native state, the PCRCR complex is highly flexible around the PfRipr tail. Binding of the non-inhibitory mAb 3F5 imposes no detectable conformational change. In contrast, inhibitory anti-EGF6-8 mAbs induce distinct conformational changes upon the tail; either by kinking (1G12, 7B3, 2H1) or extending (5B3) the EGF6-8 region that are absent in the native state (Fig. 4g). We propose that these antibody-imposed distortions restrict the natural flexibility of the PfRipr tail, locking it in conformations that compromise PCRCR complex function during invasion.

To identify additional structural features that explain the inhibitory potential of anti-Ripr mAbs, the BSA was calculated for individual EGF domains that was contributed by each anti-Ripr mAb, including 5B3 ^23^. A strong correlation was observed between EGF7 BSA and inhibitory potency (R² = 0.905), whereas neither EGF6 BSA (R² = 0.056) nor EGF8 BSA (R² = 0.556) showed a significant correlation (Fig. 4h and Supplementary Table S7). This relationship suggests that, beyond inducing conformational changes between EGF6-8 subdomains, direct engagement of EGF7 may interfere with a functionally significant role for this domain during invasion.

### PfRipr EGF6-8 is indispensable for merozoite invasion of human erythrocytes

Structural mapping identified the EGF7 subdomain as a determinant of antibody-mediated inhibition of invasion. To test directly whether EGF7 is required for PfRipr function *in situ*, a panel of rapamycin-inducible conditional *P. falciparum* lines were generated in the iGP2-DiCre background using a DiCre-loxP complementation strategy (Fig. 5a and Extended Data Fig. 9a) ^27, 31, 32^. The targeting cassette placed a loxP-flanked, recodonised wild-type *pfripr* cassette (*pfripr#1*) upstream of an out-of-frame, recodonised *pfripr#2* encoding the engineered allele with a C-terminal 3×HA tag; rapamycin-activated DiCre excises *pfripr#1* and switches *pfripr#2-ha* into frame, enabling inducible expression of the modified allele using the endogenous promoter in a strategy similar to a previous study analysing PfRh5 processing by the aspartyl protease plasmepsin X ^27^. Using this approach, parasite lines were produced encoding deletion of single EGF modules (ΔEGF6, ΔEGF7, ΔEGF8), positional swaps of adjacent modules (EGF7↔EGF6, EGF8↔EGF7) and chimeric replacements in which individual PfRipr EGF modules were substituted with heterologous EGF sequences, either PfRipr EGF3 (EGF7>EGF3) or the EGF3 subdomain from the mosquito-stage protein Pfs25 (EGF6>Pfs25, EGF7>Pfs25, EGF8>Pfs25) ^33^ (Fig. 5b). Pfs25 is a multi-EGF domain protein expressed on zygotes and ookinetes in the mosquito midgut and provides a heterologous EGF domain scaffold for distinguishing sequence-specific effects from purely structural requirements in the PfRipr tail ^34–36^.

**Fig. 5.**
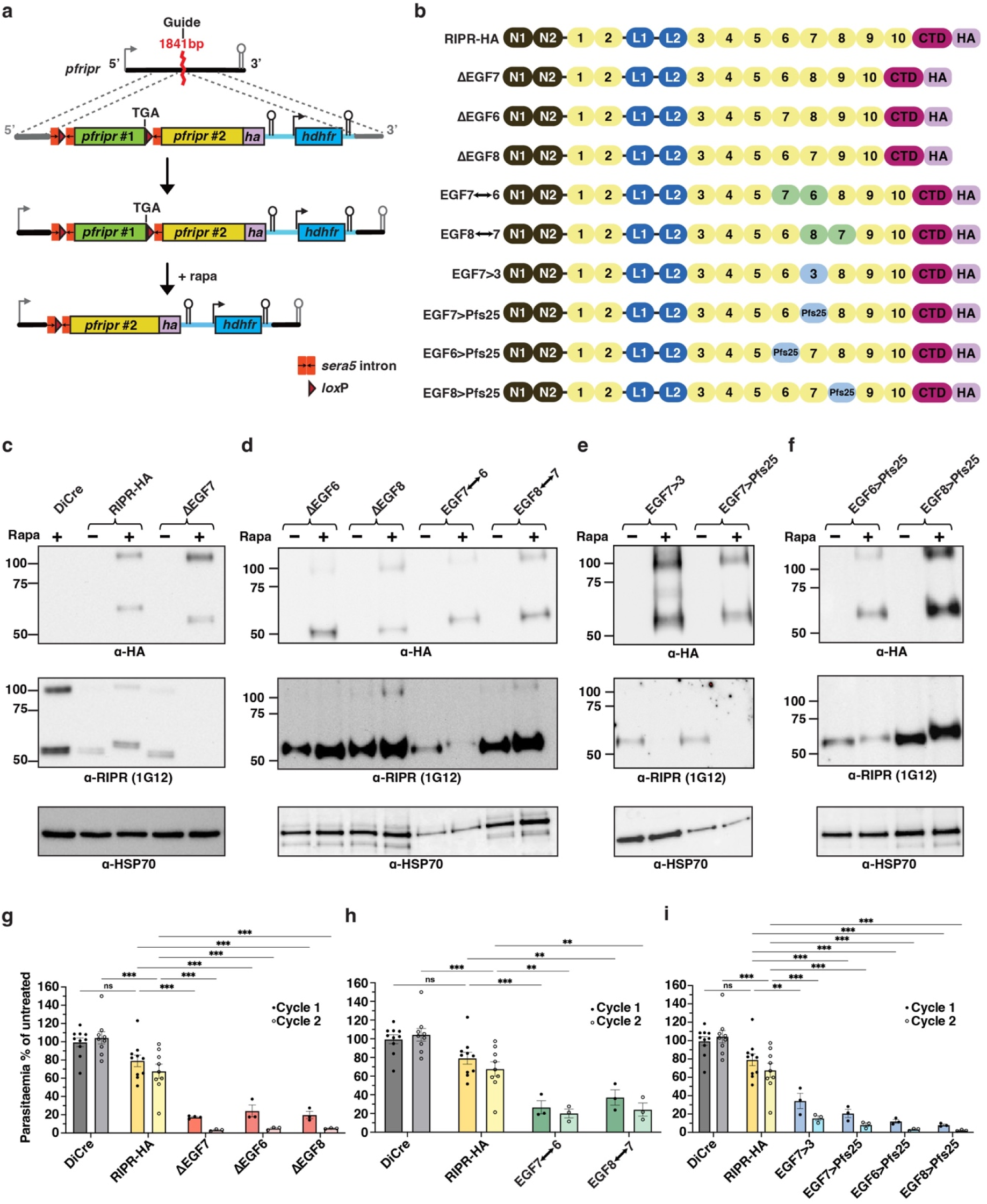
PfRipr EGF6-8 is indispensable for merozoite invasion of human erythrocytes. **a.** Schematic of the DiCre-*lox*P conditional complementation strategy for the *pfripr* gene in the *P. falciparum* iGP2-DiCre parental background line. The *pfripr* locus was targeted for construct insertion using CRISPR-Cas9 and a guide targeting position 1841 within the *pfripr* coding frame. The insert template introduces recodonised wildtype PfRipr #1 segment flanked by *lox*P sites, and an out of frame recodonised PfRipr #2 (containing a C-terminal HA tag and modifications of interest). Rapamycin-induced DiCre recombination excises PfRipr #1, activating complementation and allowing expression of the modified PfRipr #2 construct. **b.** Mutant PfRipr proteins expressed after rapamycin-induced complementation of the *pfripr* gene. From top: RIPR-HA, ΔEGF7, ΔEGF6, ΔEGF8, EGF7↔6, EGF8↔7, EGF7>3, EGF7>Pfs25, EGF6>Pfs25, EGF8>Pfs25. **c.** Validation of conditional complementation of mutant *P. falciparum* parasite lines DiCre, RIPR-HA and ΔEGF7. Schizont-stage parasites were probed with anti-HA (top), anti-PfRipr 1G12 (middle) and anti-HSP70 (bottom, loading control) antibodies. DiCre refers to the DiCre-expressing parasite line created in the iGP2 strain^32^. + and – indicates with or without rapamycin treatment. **d.** As for (c) with *P. falciparum* lines ΔEGF6, ΔEGF8, EGF7↔6 and EGF8↔7. **e.** As for (c) with *P. falciparum* lines EGF7>3, EGF7>Pfs25. **f.** As for (c) with *P. falciparum* lines EGF6>Pfs25, EGF8>Pfs25. **g-i.** Growth assays of transgenic parasites over one (black solid circles) and two (unfilled circles) replication cycles, shown as parasitaemia normalised to untreated (– rapamycin) controls of the same line at the same cycle. Three panels group transgenic lines by modification type: deletions (**g**), positional swaps (**h**) and replacements (**i**). Each panel includes the same DiCre parental line and RIPR-HA control. Except for RIPR-HA, remaining parasites in rapamycin-treated transgenic lines at cycle 2 were gametocytes; no asexual-stage parasites were detected. Cycle 1 and Cycle 2 measurements were taken from distinct samples. Error bars indicate SEM from minimum three independent experiments, as shown by the data points: N = 10 (DiCre, Cycle 1; RIPR-HA, Cycle 1); N = 9 (DiCre, Cycle 2; RIPR-HA, Cycle 2); N = 4 (ΔEGF7, Cycle 1); N = 3 (all others). Statistical significance was determined by Two-Way ANOVA with Tukey multiple comparisons test within cycles (*P* value style: not significant (ns); 0.033 (*); 0.002 (**); < 0.001 (***)).

Efficient recombination and expression of the engineered alleles following rapamycin treatment was confirmed by immunoblotting schizont-stage extracts with anti-HA, the EGF7-reactive mAb 1G12 and anti-HSP70 as a loading control (Fig. 5c-f). In parental iGP2-DiCre parasites treated with rapamycin, anti-HA produced no signal as expected whereas 1G12 detected the expected full-length PfRipr (∼120 kDa) and the PMX-processed C-terminal polypeptide (∼60 kDa) ^27^. Following rapamycin treatment, the control PfRipr-HA complementation line and each engineered line showed anti-HA bands at the predicted molecular weights, confirming efficient DiCre-mediated activation of PfRipr#2-HA expression. Domain-specific reactivity of mAb 1G12 provided independent confirmation that each modification had been introduced correctly. The *P. falciparum* lines ΔEGF7, EGF7>EGF3 and EGF7>Pfs25, in which EGF7 was absent or replaced, were mAb 1G12-negative, whereas lines retaining an intact EGF7, including ΔEGF6, ΔEGF8, EGF7↔EGF6, EGF8↔EGF7, EGF6>Pfs25 and EGF8>Pfs25, remained 1G12-reactive. The two Pfs25 replacement lines retaining EGF7 showed a detectable shift in apparent molecular weight consistent with incorporation of the heterologous domain. Collectively, the immunoblot data confirmed that DiCre-mediated recombination was highly efficient across the entire panel, with each engineered PfRipr protein expressed at the expected size consistent with the predicted domain composition.

To assess the functional consequences of each modification, growth assays were performed over two asexual replication cycles following rapamycin induction (Fig. 5g-i and Extended Data Fig. 9b). The PfRipr-HA control retained ∼70% viability relative to untreated controls across two cycles, indicating that insertion of the HA tag has some effect on PfRipr function; however, it does not block overall merozoite invasion. Comparing to this control, every engineered modification of the EGF6-8 region, that involved deletions, positional swaps and heterologous replacements, caused a profound loss of asexual blood-stage viability, only retaining ∼2%-24% viability relative to untreated controls. Flow cytometric analysis and Giemsa smear examination after two cycles revealed no detectable asexual parasites in any rapamycin-treated mutant line; the parasite forms remaining were gametocytes carried over from the starting culture. Smears of schizont-stage parasites following rapamycin treatment in the preceding ring-stage displayed morphologically normal schizont development but exhibited abnormal cellular phenotypes including an abundance of extracellular merozoites and extra-erythrocytic ring parasites, indicative of a block in the invasion capacity of the developing merozoites ^4^ (Extended Data Fig. 9c).

These genetic data demonstrated that both the primary sequence of each EGF6-8 subdomain and their precise spatial arrangement within the PfRipr tail are essential for *P. falciparum* merozoite invasion of human erythrocytes. Neither deletion nor replacement with homologous or heterologous EGF modules, nor positional swapping, was tolerated, establishing a non-redundant, sequence□ and position-specific role for the EGF6-8 module, and EGF7 in particular, in merozoite invasion and consequently parasite viability. Overall, these results are consistent with a model in which invasion-inhibitory antibodies, that include mAbs 1G12, 7B3, 2H1 and 5B3, exert their effects by engaging and functionally disabling this critical region by decreasing the flexibility of the PfRipr protein and consequently the PCRCR complex.

## Discussion

This study provides an integrated structural and genetic explanation for how antibodies targeting PfRipr disable PCRCR-dependent erythrocyte invasion by *P. falciparum* merozoites and identifies EGF7 within the PfRipr tail as a conserved, functionally indispensable vulnerability. By combining epitope mapping, kinetic characterisation, high-resolution Fab-antigen structures, combinatorial GIAs and conditional genetics, we show that neutralisation of PfRipr is determined primarily by where and how an antibody engages the tail, not simply by overall affinity or by blocking a single binary interaction within the complex.

The structural data establish two complementary antibody-mediated inhibitory mechanisms. Several potent mAbs (1G12, 7B3, 2H1) converge on EGF6-8 and make extensive contacts with EGF7, with buried surface area on this domain correlates strongly with inhibitory potency. Crystal structures further reveal that inhibitory antibodies impose structural rearrangements on the PfRipr tail: 1G12, 7B3 and 2H1 induce a pronounced kink at the EGF6-7 junction (72° versus the predicted 121° in the unbound tail), whereas 5B3 induces an extension at the EGF7-8 junction (125°), elongating the tail beyond its predicted conformation. In both cases the outcome is a constrained geometry and reduced tail flexibility that, we propose, perturbs PCRCR function at the critical moment of erythrocyte engagement. In contrast, the non-neutralising antibody 3F5 primarily engages EGF8 and does not induce any detectable angular change at the EGF7-8 junction, despite binding with high affinity and burying the largest surface area of any antibody in the panel. Together, these observations indicate that the common denominator of inhibitory activity is not a single shared conformational change but rather the restriction of natural EGF domain flexibility, whether by kinking or by locking the tail in an extended conformation, and that direct engagement of EGF7 is the principal structural correlate of potency. These structural observations extend and provide atomic-level resolution of the earlier identification of the 1G12 epitope within EGF7 ^10^, now showing that the inhibitory footprint spans EGF6-7 and that multiple structurally distinct antibodies converge on this region through divergent mechanisms that share the common outcome of restricting tail dynamics.

The suite of conditionally regulatable PfRIPR parasites provides functional validation of these structural insights. Rapamycin-inducible deletion, sequence replacement and positional swapping of EGF6-8 were uniformly lethal and blocked *P. falciparum* growth and merozoite invasion ^16, 21^, demonstrating that both the primary sequence of each subdomain and their precise spatial arrangement are non-redundant and essential for asexual blood-stage viability. The parasite’s intolerance of closely related EGF substitutions or simple domain relocations argues that EGF6-8 performs a tightly specified mechanistic role within PCRCR rather than serving as a generic structural tether. These genetic data strengthen the interpretation that neutralising antibodies disable a mechanistic lynchpin of PCRCR activity rather than merely blocking an accessory contact and explain why antibody engagement of EGF7, whether by direct surface blocking or by imposing conformational rigidity, is sufficient to inhibit merozoite invasion.

The antibody competition experiments showed that while several mAbs block PfRipr-CyRPA or PfRipr-PfPTRAMP-CSS binary interactions *in vitro*, these blocking activities did not reliably predict invasion inhibition. This was consistent with a model in which PCRCR is largely pre-assembled in micronemes prior to surface exposure ^27^, such that antibodies that merely prevent formation of a binary contact may be kinetically disadvantaged if that contact is already established before antibody access ^16^. Similarly, inhibitory potency is largely independent and uncoupled from binding kinetics, suggesting that kinetics alone cannot overcome constraints imposed by epitope location or the structural outcome of binding. Together, these data emphasise that interpreting blocking assays requires consideration of complex assembly timing, epitope accessibility and the structural consequences of antibody engagement.

The combinatorial GIA data reveal both opportunities and important caveats for immunogen design. Non-inhibitory or weakly inhibitory antibodies (3F5, 6D2) can markedly potentiate neutralisation when combined with other antibodies, including anti-Rh5 reagents, demonstrating that antibodies which alter merozoite-erythrocyte interactions without directly blocking invasion can substantially extend the window for inhibitors to act, and is analogous to the mechanism proposed for the non-inhibitory potentiating mAb R5.011 ^25, 28^. Consistent with this, 3F5, which has the largest buried surface area on EGF8, showed the strongest synergistic potential, whereas mAbs with the greatest EGF7 buried surface area (7B3, 5B3) tended to be additive or antagonistic, suggesting a possible link between EGF8 engagement and synergistic capacity. Antagonism between some inhibitory mAbs was not readily explained by steric competition, since the relevant antibodies bind non-overlapping epitopes, implying that antagonism may arise from altered complex dynamics, kinetic interference during invasion, or other properties of multi-antibody occupancy on the merozoite surface. These observations argue for pre-clinical screening of candidate combinations to identify synergistic pairings and to avoid antagonistic interactions before advancing to *in vivo* studies. More broadly, the synergy data suggest that a comprehensive map of inhibitory and potentiating epitopes across all five PCRCR components could inform the rational design of a multi-component PCRCR vaccine that deliberately combines complementary protective epitopes ^28^. Such a strategy, guided by structural definition of each inhibitory surface and empirical validation of antibody combinations, could exploit inter-complex synergy to achieve levels of invasion inhibition that exceed what any single-antigen can elicit alone.

The cross-species activity of 5B3 and the high sequence conservation across EGF6-8 in surveyed *P. falciparum* isolates is encouraging for vaccine design, indicating that EGF7-centred epitopes are conserved and capable of eliciting broadly protective responses ^23^. Nevertheless, cross-reactivity of a single mAb does not guarantee vaccine-level cross-protection, and the evolutionary capacity of *P. falciparum* means that antigenic escape requires explicit evaluation before clinical translation ^37^. Deep-mutational scanning of EGF6-8 and *in vitro* selection under mAb pressure can identify substitutions that abrogate antibody binding while preserving function, revealing escape pathways and informing immunogen designs that minimise vulnerability ^38, 39^. Parallel analysis of population sequence databases and functional testing of naturally occurring polymorphisms will clarify circulating risks and help prioritise conserved epitopes for inclusion in vaccines.

Translationally, these findings provide a blueprint for immunogen design. Stabilised EGF6-8 constructs that present the protective EGF7 surface in its native spatial context should be prioritised over isolated EGF7 fragments, since our genetic data show that adjacent domains and their relative positions are critical for both function and native epitope geometry. Multiepitope or multivalent display formats are attractive because EGF6-8 contains multiple independent inhibitory epitopes; coupling EGF7-focused immunogens with components that elicit potentiating specificities (EGF8 or selected core epitopes) and with validated targets such as PfRh5 may amplify potency and breadth. Concurrent higher-order structural characterisation of the intact PCRCR complex, escape-mapping experiments and *in vivo* efficacy studies with immunogen optimisation would accelerate translation.

In conclusion, our integrated structural, biophysical and genetic dissection identifies EGF7 within PfRipr as a conserved, functionally essential element whose direct engagement or antibody-induced distortion potently neutralises merozoite invasion. These mechanistic insights reconcile and extend prior mapping studies ^10, 25^ and refine our understanding of PfRipr within the PCRCR assembly. Taken together with the earlier identification of the 1G12 inhibitory epitope in EGF7 ^10^ and preclinical immunogenicity data demonstrating that PfRipr alone can elicit more potent invasion-inhibitory antibodies than PfRh5 or PfCyRPA individually, and that a triple-antigen RCR combination offers little improvement over PfRh5 alone ^10, 26^, our structural data converge on a single dominant inhibitory epitope centred on EGF7 with critical contacts extending into EGF6 and 8. This epitope is the convergent target of the most potent inhibitory anti-PfRipr antibodies characterised to date, and its precise geometry spanning the EGF6-7 junction is essential for productive antibody engagement. The lack of improvement seen with the triple RCR combination in preclinical assays most likely reflects immunodominance of the PfRipr core: when whole-antigen combinations are used, the antibody response is dominated by PfRipr, and specifically by the core, such that the addition of PfCyRPA and PfRh5 antigens does not significantly augment the overall inhibitory titre ^25, 26^. This interpretation is consistent with our synergy data, which show that potentiation by non-inhibitory antibodies targeting EGF8 or the PfRipr core requires deliberate co-administration at defined concentrations rather than arising spontaneously from a polyclonal response to a mixed antigen preparation. Together, these observations suggest that realising the full synergistic potential of multi-component PCRCR vaccines will require structure-guided immunogen design that focuses the response on the most protective epitopes, rather than simply combining whole antigens, and that empirical optimisation of antigen combinations and delivery will be essential to translate the synergy observed here into durable, broadly protective immunity.

## Acknowledgements

The authors thank Australian Red Cross Lifeblood for the supply of human red blood cells and serum. We thank G. Kong from the Monash Macromolecular Crystallisation Facility (MMCF, Clayton, VIC, Australia) for assistance with setting up crystallisation screens (https://www.monash.edu/researchinfrastructure/mmcp), and the Bio21-WEHI Crystallisation Facility within The Bio21 Molecular Science and Biotechnology Institute, University of Melbourne, for crystallisation support. We acknowledge the Bio21 Advanced Microscopy Facility for electron microscopy infrastructure, the WEHI Cryo-EM Facility for cryo-electron microscopy data collection, and the WEHI Antibody Facility and WEHI Protein Production Facility for production of monoclonal antibodies and recombinant proteins. This research was undertaken in part using the MX2 beamline at the Australian Synchrotron, part of the Australian Nuclear Science and Technology Organisation (ANSTO) and made use of the Australian Cancer Research Foundation (ACRF) detector. We thank Holger Kanzler, Jacqueline Kirchner, Matthias Paulsner and Annie Zumsteg from the Gates Foundation for helpful discussion. This work was supported by the Gates Foundation (INV-074041), the National Health and Medical Research Council of Australia (NHMRC; grants GNT1173049, GNT1194535, GNT2025925, GNT2034643, GNT 2016827 and GNT2045875), the Drakensberg Trust and infrastructure support by the Victorian State Government Operational Infrastructure Support grant.

## Author Contributions

XX, NCJ and SWS expressed proteins, and performed and analysed biophysical experiments. XX solved crystal structures with assistance from SWS. XX and TCM performed mass photometry analysis. SS and XX analysed cryo-EM data. QZ and XX prepared cryo-EM grids and AL carried out cryo-EM data collection. XX and PSL performed and analysed parasite growth inhibition assays. XX, QZ, TCM and RC performed and analysed pairwise combinatorial synergy assays. DSM, XX, MWAD and SL designed the transfection constructs. XX and DSM designed and analysed all transgenic parasites. MYN and AEB performed antigen sequence conservation analysis. XX, AFC and SWS designed and interpreted experiments. XX, AFC and SWS wrote the manuscript. All authors read and edited the manuscript.

## Materials and Methods

### Recombinant protein expression

All gene sequences used were retrieved from the VEuPathDB (accessed through www.plasmodb.org) ^41^ from reference strains (3D7 for *P. falciparum*). All genes were synthesized by Genscript (Singapore) unless otherwise stated.

### *P. falciparum* constructs and protein expression

Recombinant PfPTRAMP-CSS for competition studies was produced as previously described. Briefly, PfPTRAMP, comprising residues 31 to 307, was engineered to remove four potential N-linked glycan sites by mutation: Asn112, Asn149, Asn155 to Gln, and Thr197 to Ala to eliminate glycosylation at Asn195. Full length PfCSS was similarly engineered to remove all six predicted N-linked glycosylation sites by mutating Ser76, Thr90, Ser194, Thr236 and Thr263 to Ala and Asn283 to Gln. PfPTRAMP and PfCSS were co-expressed in Sf21 cells and purified via C-tagXL affinity matrix chromatography (Thermo Fisher Scientific) ^23^. Recombinant full-length PfRipr (20-1086) with C-terminal His tag, PfCyRPA, PfRh5 constructs were produced as previously described ^16, 23^. Full-length PfRipr with C-terminal Strep-tag II was purified from supernatant via binding and eluting using a Strep-Tactin XT Sepharose chromatography resin following methods previously described for PvRipr_22–1074_Strep ^22, 23^.

PfRipr^tail^(717-1086), PfRipr^EGF9-10-CTD^(899-1086) and PfRipr^CTD^(981-1086), were all synthesised by Genscript and purified in an identical fashion to full-length PfRipr with C-terminal His-tag. PfRipr^EGF6-8^ (771-897), PfRipr^EGF6-7^ (771-854), PfRipr^EGF6^ (771-814) and PfRipr^EGF8^ (858-897) were synthesized and subcloned into pET28a vector each yielding a construct with an N-terminal 6x His tag followed by a Tobacco Etch Virus (TEV) site and produced in an identical fashion as PvRipr^EGF6-8^ as previously described ^23^. PfRipr^EGF7-8^ (818-897) was synthesized and subcloned into pcDNA3.4 vector with the addition of a N-terminal 6xHis tag followed by a TEV site and produced in an identical fashion as PvRiprEGF^7–8^ as previously described ^23^.

The stabilised PfPTRAMP-CSS-Ripr (PCR) co-complex was engineered by introducing a disulfide-linkage between PfPTRAMP and PfRipr. Briefly, an AlphaFold3 prediction of the PfPTRAMP-CSS-Ripr^tail^ complex were used as input for Disulfide by Design 2.0, to identify residue pairs for cysteine substitution ^42^. Residue Asn195 from PfPTRAMP and residue Lys1024 from PfRipr were selected and mutated to cysteine to enable disulfide bond formation. These constructs were combined with PfCSS, which was engineered to remove all six predicted N-linked glycosylation sites by mutating Ser76, Ser194, Thr236 and Thr263 to Ala, and Asn88 and Asn283 to Gln. The complex was then co-expressed in Sf21 cells and purified via Ni-NTA agarose resin (Qiagen) chromatography, followed by C-tagXL affinity matrix chromatography (Thermo Fisher Scientific) and size exclusion chromatography (S200 Increase 10/300 GL, Cytiva).

### Antibody and Fab production

Anti-PfRipr monoclonal antibodies were raised in mice as per the Walter and Eliza Hall Animal Ethics Committee approved procedures. All monoclonal antibodies were produced by the WEHI Antibody Facility. Mice were immunized three times with 10 μg of full-length PfRipr, followed by a booster immunization with 20 μg prior to hybridoma fusion. Following initial hybridoma cloning, culture supernatants were screened for PfRipr binding and epitope competition using ELISA and BLI. Based on these analyses, nine PfRipr-specific hybridomas were selected (Fig. 1a and d) for final cloning and antibody expression. Anti-PfRipr hybridomas were sequenced as previously described ^43^.

Fabs for 1G12, 7B3, 2H1 and 3F5 were generated by papain digestion of IgG and purified by Protein G affinity chromatography (HiTrap Protein G, Cytiva), followed by size-exclusion chromatography (S200 Increase 10/300 GL, Cytiva). In addition, for 7B3, 2H1, 3F5 and 6D2, the *IGH*, *IGK* or *IGL* variable regions were synthesized and cloned into pcDNA3.4 TOPO expression vectors upstream of the appropriate human *IGK*, *IGL* or *CH1* constant regions for recombinant Fab expression in expi293F cells (Thermo Fisher Scientific). Fabs were purified from supernatant via HiTrap MabSelect VL, HiTrap Protein G column or HiTrap LambdaFabSelect as appropriate, followed by size exclusion chromatography (S200 Increase 10/300 GL, Cytiva).

The *IGH*, *IGK* and *IGL* variable regions for anti-Rh5 Abs R5.011 and R5.016 were synthesized and subcloned into pcDNA3.4 TOPO expression vectors upstream of the appropriate human *IGK*, *IGL* or *CH1-Fc* constant regions for recombinant hIgG1 expression in expi293F cells (Thermo Fisher Scientific). Antibodies were purified from supernatant via 5mL HiTrap Prism A column (Cytiva), followed by size exclusion chromatography (S200 Increase 10/300 GL, Cytiva).

### Biolayer interferometry (BLI)

BLI experiments were carried out on an Octet Red96e (Sartorius) at 25°C. For antibody kinetics, anti-Mouse IgG Fc Capture (AMC) biosensors (Sartorius) were used to immobilise mouse monoclonal antibodies at a concentration of 10 μg/mL in kinetics buffer (PBS, pH 7.4, 0.1% (w/v) BSA, 0.02% (v/v) Tween-20). Biosensors were initially dipped in kinetics buffer for 30 seconds to establish a baseline signal, and then dipped into wells containing the antibodies, followed by another 60 second baseline. After the second baseline step, the ligands were then dipped into wells containing two-fold dilution series of analyte (PfRipr constructs). Association was measured for 120 seconds and then the biosensors were dipped into kinetics buffer to measure the dissociation for another 120 seconds. Data were analysed using Sartorius’ Data Analysis software 11.0. Kinetic curves were fitted using a 1:1 binding model.

Competition studies for anti-PfRipr mAbs were performed using Nickel-Nitrilotriacetic acid (Ni-NTA) biosensors (Sartorius). For epitope binning assays, biosensors were first dipped into wells containing kinetics buffer for 30 seconds, then dipped into wells containing 20 µg/mL of full-length PfRipr with C-terminal His tag for 120 seconds, followed by another 30 second baseline step. Sensors were then dipped into wells containing 20 µg/mL of primary mAbs for 300 seconds. A final baseline step was performed before the biosensors were dipped into secondary mAb (20 μg/mL) to assess competition. For PfPTRAMP-CSS and PfCyRPA competition studies, biosensors were first dipped into wells containing kinetics buffer for 30 seconds, then dipped into wells containing 20 µg/mL of PfRipr_20-1086_2xA for 120 seconds, followed by another 30 second baseline step. Sensors were then dipped into wells containing 2 µM of PfPTRAMP-CSS or 50 µg/mL of primary mAbs diluted in kinetics buffer, for 300 seconds. A final baseline step was performed before the biosensors were dipped into mAb (50 μg/mL), PfPTRAMP-CSS (2 µM) or PfCyRPA-Rh5 premix (2 μM) to assess competition. Data were analysed using Sartorius’ Data Analysis software 11.0 and the epitope bins assessed by normalisation and manual curation.

### *In vitro* Growth inhibition assays (GIAs)

To measure the *in vitro* growth inhibitory activity of the anti-PfRipr mAbs, standard growth inhibition assays were performed as described previously ^10, 16^.

Combinatorial GIAs were performed as previously described ^44^ and the predicted Bliss additivity was calculated using the measured activity of each antibody independently using formulas previously described ^29^. One mAb was titrated across a four-fold seven-step dilution series starting from 2 mg/mL, while a second mAb was held at a constant concentration to achieve approximately 30% GIA. For combinations involving 3F5 and R5.011, these Abs were instead held at 500 µg/mL (not 30% GIA) as neither Ab exhibits GIA activity when tested alone. After incubation for 48 hr, each well was fixed at room temperature for 30 min with 50 µL of 0.25% glutaraldehyde (ProSciTech) diluted in human tonicity PBS. Following centrifugation at 1,200 rpm for 2 min, supernatants were discarded, parasites were resuspended in 50 µL PBS and stained with 50 µL SYBR Green (Invitrogen) diluted in PBS at 37 °C in the dark. The parasitaemia of each well was determined by counting 200,000 cells by flow cytometry using a NovoCyte Flow Cytometer (Agilent Technologies).

### Protein crystallization

Fabs 1G12 and 7B3 were co-complexed with PfRipr^EGF6-8^, 2H1 with PfRipr^EGF6-7^ and 3F5 Fab with PfRipr^EGF7-8^ at a 1:2 Fab:antigen molar ratio. Excess antigen was removed by size-exclusion chromatography (S200 Increase GL 10/300, Cytiva) in HBS buffer, pH 7.5. Complexes were then concentrated to 5 mg/ml and mixed 1:1 with mother liquor and set up in hanging or sitting drop crystallization experiments. 2H1-PfRipr^EGF6-7^ crystals grew in 0.2 M LiSO_4_, 30% PEG3350, 0.1 M Tris pH 9, and were cryoprotected in 15% glycerol; 3F5-PfRipr^EGF7-8^ crystals grew in 0.2 M LiSO_4_, 20% PEG3350, 0.1 M Tris pH 8, and were cryoprotected in 15% ethylene glycol. 1G12-PfRipr^EGF6-8^ crystals grew in 2 M (NH_4_)_2_SO_4_, 0.1 M Bis-Tris pH 5.5, and 7B3-PfRipr^EGF6-8^ crystals grew in 0.2 M (NH_4_)_2_SO_4_, 20% (w/v) PEGSB, 10% (w/v) ethylene glycol, 0.01 M cadmium chloride hydrate, 0.1 M PIPES pH 6.5. Both crystals were cryoprotected in 20 % (w/v) ethylene glycol. Diffraction data were collected with the MX2 beamline at the Australian Synchrotron (Clayton, Australia) at 100 K (λ = 0.9537 Å). Statistics are in Supplementary Table S2.

### Structure determination and model building

All diffraction data were processed and reduced with the XDS package ^45^ before being scaled and merged using Aimless ^46^ in the CCP4 suite ^47^. The program Matthews ^48^ was used to estimate the number of molecules in the asymmetric unit. 1 molecule was estimated in the asymmetric unit for 1G12-PfRipr^EGF6-8^ and 7B3-PfRipr^EGF6-8^; 2 molecules for 2H1-PfRipr^EGF6-7^ and 4 molecules for 3F5-PfRipr^EGF7-8^. The AlphaFold3 model of the PfRipr antigens and the crystal structure of an HIV-2 neutralizing Fab fragment (PDB: 3NZ8) ^49^ with the CDRs deleted, as well as the crystal structure of 5B3-PvRipr^EGF7-8^ (PDB: 9YIO) ^23^ were used as models for molecular replacement using Phaser ^50^. Alternating rounds of structure building and refinement were carried out using Phenix ^51^ and assessed and modified with Coot ^52^. Refinement and model statistics are found in Supplementary Table S2. Interactions between PfRipr antigens and Fabs were analyzed using PISA server (https://www.ebi.ac.uk/pdbe/pisa/) and CCP4 Contact ^47^ and are summarized in Supplementary Table S3-S6. All Fabs were renumbered according to the Kabat numbering system, as determined by ANARCI ^53^.

### Structure prediction

Prediction of PCRCR complexes from *P. falciparum* was performed using the AlphaFold3 server ^30^.

### Stabilised PfPCRCR–Fabs co-complexation

Anti-PfRipr 7B3, 2H1, 3F5 and 6D2 recombinant Fabs, in addition to recombinant PfCyRPA and PfRh5 were co-complexed with the stabilised PfPTRAMP-CSS-Ripr (PCR) complex to form the stabilised PfPCRCR-Fabs co-complex. The ratio of Fabs: CyRPA:PfRh5:PCR pre-incubated was 1.5:2:2:1. Excess antigens and Fabs were removed by size-exclusion chromatography (Superose 6 Increase GL 10/300, Cytiva) in HBS buffer, pH 7.5, with peak fractions assessed via SDS-PAGE and pooled.

### Mass photometry

Mass photometry experiments were carried out on a Two^MP^ mass photometer (Refeyn). Gasket wells were focused after the addition of 10 μL of filtered 20mM HEPES buffer pH 7.5. Once focused, 10 μL of stabilised PfPCRCR-Fabs co-complex sample at 10 nM was added, mixed, and events were recorded for one minute using AcquireMP (Refeyn). Raw data processing and representative histograms were generated using DiscoverMP (Refeyn). Molecular weight of PfPCRCR-Fabs co-complex was estimated using a standard curve generated from recombinant human glutamine synthetase (Cedarlane). Molecule counts were used to determine levels of protein complexes.

### Cryo-Electron microscopy sample preparation and data acquisition

A 3.5 µl freshly purified PfPCRCR–Fabs co-complex at 0.2 mg/mL was applied to plasma-cleaned UltrAu foil grids (300 mesh, 1.2/1.3) and then blotted for 3.5 seconds with a blot force of 0 before being plunged into liquid ethane on an FEI Vitrobot Mark IV (Thermo Fisher Scientific) operating at 4°C and 100% humidity. The grids were imaged on a Titan Krios G4 microscope at the Ian Holmes Imaging Centre, Bio21, University of Melbourne, operating at 300kV on a Falcon IV detector. A total of 10,000 movies were collected at a pixel size of 0.639 Å/pixel, with a total dose of 40 e-/Å2. Images were acquired with a defocus range of −0.5 to −1.6 μm.

### Data processing

CryoSPARC v4.6.2 ^54^ was used for all image processing steps except particle picking. Beam-induced motion correction was performed using Patch Motion Correction, followed by estimation of contrast transfer function (CTF) parameters using Patch CTF Estimation. Low-quality micrographs were removed using the Manually Curate Exposures job based on a maximum CTF fit resolution cutoff of 5 Å, resulting in 9,149 micrographs for downstream processing.

Particles were automatically picked using PartiNet, a dynamically adaptive AI-based particle picker, on denoised micrographs with a confidence threshold of 0.1, yielding 1,885,614 initial particle coordinates ^55^. PartiNet processing was performed according to the published protocol and associated workflow documentation (https://wehi-researchcomputing.github.io/PartiNet/). Extracted particles were subjected to three rounds of reference-free 2D classification to remove junk particles and poorly aligned classes. Classification was performed using 100 classes, an uncertainty factor of 1, 20 final full iterations, 80 online-EM iterations, and a batch size of 400 particles.

Following 2D classification, 572,751 particles were selected for ab initio reconstruction into three classes. The best-resolved class was refined through three rounds of heterogeneous refinement and non-uniform refinement. The final round of non-uniform refinement yielded a 3.50 Å reconstruction but exhibited severe preferred orientation. To alleviate orientation bias, orientation rebalancing was performed using the random assignment mode at the 80th percentile, resulting in a subset of 68,128 particles. These particles were subjected to an additional round of non-uniform refinement, producing a final reconstruction at 5.14 Å resolution.

The handedness of the final cryo-EM map (EMDB-80770) was corrected, and used for rigid-body fitting of the 7B3, 2H1, and 3F5 Fab-PfRipr crystal structures (PDB IDs: 24HV, 24HW, 24HY) together with AlphaFold3-predicted models of the EGF5 and EGF9 subdomains in UCSF ChimeraX. Refinement and model statistics are found in Supplementary Table S9.

### Data visualisation

All data visualisation, centre of mass and angle between axes determination were done in University of California, San Francisco (UCSF) ChimeraX versions 1.8-1.11 (https://www.cgl.ucsf.edu/chimerax/) ^56^. PyMOL was utilised for structure alignment and calculation of root mean square deviation (RMSD) (https://www.pymol.org/) ^57^.

### Assembly of plasmid constructs for mutation of PfRIPR

For the guide and Cas9 containing vector targeting *pfripr*, oligos DM945/DM946 (Integrated DNA Technologies) were designed to induce a double-stranded break at position 1841 of the *pfripr* coding sequence and cloned using InFusion into pUF1-Cas9G using previously published methods ^21^. This vector was used as the guide vector for all *pfripr* modifications.

The initial homology-directed repair (HDR) construct used for all *pfripr* modifications was designated RIPR-HA. All gene synthesis and cloning were performed by GenScript Biotech.

A 625 bp 3′ homology region of *pfripr* was synthesised and cloned into p1.2-RBX1-HA ^58, 59^ using *Eco*RI and *Kas*I restriction sites, generating p1.2-RBX1-HA-PfRIPR3′. Subsequently, a 591 bp 5′ homology region together with two sequential, differentially codon-optimised *pfripr* coding sequences were synthesised and inserted into p1.2-RBX1-HA-PfRIPR3′ via *Not*I and *Xma*I restriction sites to generate the final RIPR-HA construct. The first codon-optimised *pfripr* sequence was flanked by *Lox*P sites embedded within heterologous *sera5* intron sequences. The second codon-optimised *pfripr* sequence was positioned downstream, inserted between *Spe*I and *Xma*I restriction sites, and maintained out of frame relative to the upstream coding sequence. In the RIPR-HA construct, the first and second codon-optimised *pfripr* sequences were identical except that the second sequence encoded a C-terminal 3×HA epitope tag. For all subsequent *pfripr* modifications, RIPR-HA was used as the parental backbone. Newly synthesised *pfripr* variants were introduced by replacing the second codon-optimised sequence within the *Spe*I/*Xma*I sites (Fig. 5a).

### Parasite transfection and recovery of transgenic parasites

A combined linearized HDR plasmid (100 µg) and circular guide plasmid (100 µg) were resuspended in 50 µl Tris-EDTA buffer (Sigma) and transfected into synchronised schizonts suspended in 85 µl Solution 1 and 15 µl Solution 2 of Basic Parasite Nucleofector Kit 2 (Lonza). Program U-33 with the Amaxa Nucleofector 2b (Lonza) was used for transfection. The marker free iGP2-DiCre parasite line was generated by introducing the pBSPfs47DiCre vector (a gift from Ellen Knuepfer ^31^) containing homology regions to the non-essential *pfs47* locus along with the pDC2-BSD-Cas9-Pfs47 Cas9 expressing guide plasmid ^59^. Following Nucleofection the parasites were selected on Blasticidin S HCL (2.5 μg/mL) for 5 days. Drug pressure was then removed to permit loss of circular plasmid and transgenic parasites were detected 14 days later, at which stage clonal iGP2-DiCre parasites were cloned by limiting dilution to obtain clonal positive parasites. For all transfections designed to introduce *pfripr* modifications, parasites with an integrated human dihydrofolate reductase gene (*hdhfr*) cassette were selected and maintained on 2.5 nM WR99210 (Jacobus Pharmaceuticals).

### Parasite growth assay

iGP2-DiCre and transgenic ring stage parasites were sorbitol-synchronised following protocols previously described ^27^ then diluted to approximately 0.5% and 0.1% parasitaemia for one-cycle and two-cycle growth respectively, with or without 10 nM rapamycin treatment. After approximately 72 and 120 hours, parasite smears were taken for Giemsa staining, and parasites were stained with Hoechst33342 (Invitrogen) diluted in Roswell Park Memorial Institute (RPMI) 1640 medium supplemented with 25 mM 4-(2-hydroxyethyl)-1-piperazineethanesulfonic acid (HEPES) at 37 °C in the dark. Parasitaemia was determined by counting 200,000 cells by flow cytometry using a NovoCyte Penteon Flow Cytometer (Agilent Technologies).

### SDS-PAGE and immunoblotting

iGP2-DiCre parental and transgenic ring-stage parasites were sorbitol-synchronised and treated with or without 10 nM rapamycin. *P. falciparum* parasites were grown to schizont-stage, parasite smears taken for Giemsa staining, and infected erythrocytes lysed with 0.15% saponin (w/v). Schizont pellets were further suspended in non-reducing SDS sample buffer, boiled at 95 °C for 10 minutes, and resolved on 4-12% NuPAGE Bis-Tris polyacrylamide gels (Invitrogen) followed by transfer to nitrocellulose membranes by electroblotting. Membranes were blocked overnight with 5% Skim Milk in PBS + 0.1% Tween-20, then probed with the following antibodies: HRP-conjugated anti-HA (1:1,000; Roche); anti-PfRipr 1G12 primary (1:2,000) followed by HRP-conjugated anti-mouse secondary (1:2,000; Sigma); and anti-HSP70 ^60^ primary (1:20,000) followed by HRP-conjugated anti-rabbit secondary (1:4,000; Merck) as a loading control. Bands were visualized using SuperSignal West Pico PLUS chemiluminescent substrate (Thermo Fisher Scientific) and imaged on a ChemiDoc Imaging System (Bio-rad).

## References

1. Weiss, D.J. et al. Mapping the global prevalence, incidence, and mortality of *Plasmodium falciparum*, 2000-17: a spatial and temporal modelling study. Lancet 394, 322–331 (2019).

2. Battle, K.E. et al. Mapping the global endemicity and clinical burden of *Plasmodium vivax*, 2000-17: a spatial and temporal modelling study. Lancet 394, 332–343 (2019).

3. Cowman, A.F., Tonkin, C.J., Tham, W.H. & Duraisingh, M.T. The Molecular Basis of Erythrocyte Invasion by Malaria Parasites. Cell Host Microbe 22, 232–245 (2017).

4. Weiss, G.E. et al. Revealing the Sequence and Resulting Cellular Morphology of Receptor-Ligand Interactions during *Plasmodium falciparum* Invasion of Erythrocytes. PLoS Pathog. 11, e1004670 (2015).

5. Hayton, K. et al. Erythrocyte binding protein PfRH5 polymorphisms determine species-specific pathways of *Plasmodium falciparum* invasion. Cell Host Microbe 4, 40–51 (2008).

6. Baum, J. et al. Reticulocyte-binding protein homologue 5 -an essential adhesin involved in invasion of human erythrocytes by *Plasmodium falciparum*. Int. J. Parasitol. 39, 371–373 (2009).

7. Douglas, A.D. et al. The blood-stage malaria antigen PfRH5 is susceptible to vaccine-inducible cross-strain neutralizing antibody. Nat. Commun. 2, 601 (2011).

8. Bustamante, L.Y. et al. A full-length recombinant *Plasmodium falciparum* PfRH5 protein induces inhibitory antibodies that are effective across common PfRH5 genetic variants. Vaccine 31, 373–379 (2013).

9. Douglas, A.D. et al. Neutralization of *Plasmodium falciparum* merozoites by antibodies against PfRH5. J. Immunol. 192, 245–258 (2014).

10. Healer, J. et al. Neutralising antibodies block the function of Rh5/Ripr/CyRPA complex during invasion of *Plasmodium falciparum* into human erythrocytes. Cell Microbiol. 21, e13030 (2019).

11. Minassian, A.M. et al. Reduced blood-stage malaria growth and immune correlates in humans following RH5 vaccination. Med (N Y) 2, 701–719 e719 (2021).

12. Silk, S.E. et al. Blood-stage malaria vaccine candidate RH5.1/Matrix-M in healthy Tanzanian adults and children; an open-label, non-randomised, first-in-human, single-centre, phase 1b trial. Lancet Infect Dis 24, 1105–1117 (2024).

13. Richards, J.S. et al. Identification and Prioritization of Merozoite Antigens as Targets of Protective Human Immunity to *Plasmodium falciparum* Malaria for Vaccine and Biomarker Development. J. Immunol. 191, 795–809 (2013).

14. Conway, D.J. Paths to a malaria vaccine illuminated by parasite genomics. Trends Genet 31, 97–107 (2015).

15. Lopaticki, S. et al. Reticulocyte and erythrocyte binding-like proteins function cooperatively in invasion of human erythrocytes by malaria parasites. Infect Immun 79, 1107–1117 (2011).

16. Scally, S.W. et al. PCRCR complex is essential for invasion of human erythrocytes by *Plasmodium falciparum*. Nat Microbiol 7, 2039–2053 (2022).

17. Chen, L. et al. An EGF-like protein forms a complex with PfRh5 and is required for invasion of human erythrocytes by *Plasmodium falciparum*. PLoS Pathog. 7, e1002199 (2011).

18. Favuzza, P. et al. Structure of the malaria vaccine candidate antigen CyRPA and its complex with a parasite invasion-inhibitory antibody. Elife 6 (2017).

19. Chen, L. et al. Structural basis for inhibition of erythrocyte invasion by antibodies to *Plasmodium falciparum* protein CyRPA. Elife 6 (2017).

20. Wright, K.E. et al. Structure of malaria invasion protein RH5 with erythrocyte basigin and blocking antibodies. Nature 515, 427–430 (2014).

21. Volz, J.C. et al. Essential Role of the PfRh5/PfRipr/CyRPA Complex during *Plasmodium falciparum* Invasion of Erythrocytes. Cell Host Microbe 20, 60–71 (2016).

22. Wong, W. et al. Structure of *Plasmodium falciparum* Rh5-CyRPA-Ripr invasion complex. Nature 565, 118–121 (2019).

23. Seager, B.A. et al. PTRAMP, CSS and Ripr form a conserved complex required for merozoite invasion of Plasmodium species into erythrocytes. Nat Commun 17, 1780 (2026).

24. Farrell, B. et al. The PfRCR complex bridges malaria parasite and erythrocyte during invasion. Nature 625, 578–584 (2024).

25. Williams, B.G. et al. Development of an improved blood-stage malaria vaccine targeting the essential RH5-CyRPA-RIPR invasion complex. Nat Commun 15, 4857 (2024).

26. Healer, J. et al. RH5.1-CyRPA-Ripr antigen combination vaccine shows little improvement over RH5.1 in a preclinical setting. Front Cell Infect Microbiol 12, 1049065 (2022).

27. Triglia, T. et al. Plasmepsin X activates the PCRCR complex of *Plasmodium falciparum* by processing PfRh5 for erythrocyte invasion. Nat Commun 14, 2219 (2023).

28. Alanine, D.G.W. et al. Human Antibodies that Slow Erythrocyte Invasion Potentiate Malaria-Neutralizing Antibodies. Cell 178, 216–228 e221 (2019).

29. Azasi, Y. et al. Bliss’ and Loewe’s additive and synergistic effects in *Plasmodium falciparum* growth inhibition by AMA1-RON2L, RH5, RIPR and CyRPA antibody combinations. Sci. Rep. 10, 11802 (2020).

30. Abramson, J. et al. Accurate structure prediction of biomolecular interactions with AlphaFold 3. Nature 630, 493–500 (2024).

31. Knuepfer, E., Napiorkowska, M., van Ooij, C. & Holder, A.A. Generating conditional gene knockouts in *Plasmodium* -a toolkit to produce stable DiCre recombinase-expressing parasite lines using CRISPR/Cas9. Sci. Rep. 7, 3881 (2017).

32. Boltryk, S.D. et al. CRISPR/Cas9-engineered inducible gametocyte producer lines as a valuable tool for *Plasmodium falciparum* malaria transmission research. Nat Commun 12, 4806 (2021).

33. Scally, S.W. et al. Molecular definition of multiple sites of antibody inhibition of malaria transmission-blocking vaccine antigen Pfs25. Nat Commun 8, 1568 (2017).

34. Vermeulen, A.N., Roeffen, W.F., Henderik, J.B., Ponnudurai, T. & Beckers *Plasmodium falciparum* transmission blocking monoclonal antibodies recognize monovalently expressed epitopes. Dev. Biol. Stand. 62, 91–97 (1985).

35. Vermeulen, A.N., van, D.J., Brakenhoff, R.H. & Lensen, T.H. Characterization of *Plasmodium falciparum* sexual stage antigens and their biosynthesis in synchronised gametocyte cultures. Mol. Biochem. Parasitol. 20, 155–163 (1986).

36. Saxena, A.K., Wu, Y. & Garboczi, D.N. *Plasmodium* p25 and p28 surface proteins: potential transmission-blocking vaccines. Eukaryot Cell 6, 1260–1265 (2007).

37. Rawlinson, T.A. et al. Structural basis for inhibition of *Plasmodium vivax* invasion by a broadly neutralizing vaccine-induced human antibody. Nat Microbiol 4, 1497–1507 (2019).

38. Fowler, D.M. & Fields, S. Deep mutational scanning: a new style of protein science. Nat Methods 11, 801–807 (2014).

39. Greaney, A.J. et al. Complete Mapping of Mutations to the SARS-CoV-2 Spike Receptor-Binding Domain that Escape Antibody Recognition. Cell Host Microbe 29, 44–57 e49 (2021).

40. Krissinel, E. & Henrick, K. Inference of macromolecular assemblies from crystalline state. J Mol Biol 372, 774–797 (2007).

41. Amos, B. et al. VEuPathDB: the eukaryotic pathogen, vector and host bioinformatics resource center. Nucleic Acids Res 50, D898–D911 (2022).

42. Craig, D.B. & Dombkowski, A.A. Disulfide by Design 2.0: a web-based tool for disulfide engineering in proteins. BMC Bioinformatics 14, 346 (2013).

43. Subas Satish, H.P., et al. NAb-seq: an accurate, rapid, and cost-effective method for antibody long-read sequencing in hybridoma cell lines and single B cells. MAbs 14, 2106621 (2022).

44. Williams, A.R. et al. Enhancing blockade of *Plasmodium falciparum* erythrocyte invasion: assessing combinations of antibodies against PfRH5 and other merozoite antigens. PLoS Pathog. 8, e1002991 (2012).

45. Kabsch, W. XDS. Acta. Crystallogr. D Biol. Crystallogr. 66, 125–132 (2010).

46. Evans, P.R. & Murshudov, G.N. How good are my data and what is the resolution? Acta Crystallogr. D. Biol. Crystallogr. 69, 1204–1214 (2013).

47. Collaborative Computational Project, N. The CCP4 Suite: Programs for Protein Crystallography. Acta Cryst. D. Biol. Crystallogr. 50, 760–763 (1994).

48. Kantardjieff, K.A. & Rupp, B. Matthews coefficient probabilities: Improved estimates for unit cell contents of proteins, DNA, and protein-nucleic acid complex crystals. Protein Sci 12, 1865–1871 (2003).

49. Uchtenhagen, H. et al. Crystal structure of the HIV-2 neutralizing Fab fragment 7C8 with high specificity to the V3 region of gp125. PLoS One 6, e18767 (2011).

50. McCoy, A.J. et al. Phaser crystallographic software. J Appl Crystallogr 40, 658–674 (2007).

51. Adams, P.D. et al. PHENIX: a comprehensive Python-based system for macromolecular structure solution. Acta Cryst. D. Biol. Crystallogr. 66, 213–221 (2010).

52. Emsley, P. & Cowtan, K. Coot: model-building tools for molecular graphics. Acta Cryst. D. Biol. Crystallogr. 60, 2126–2132 (2004).

53. Dunbar, J. & Deane, C.M. ANARCI: antigen receptor numbering and receptor classification. Bioinformatics 32, 298–300 (2016).

54. Punjani, A., Rubinstein, J.L., Fleet, D.J. & Brubaker, M.A. cryoSPARC: algorithms for rapid unsupervised cryo-EM structure determination. Nat Methods 14, 290–296 (2017).

55. Perera, M., et al. PartiNet is a dynamic adaptive neural network for high-performance particle picking in cryo-electron microscopy. bioRxiv (2026).

56. Pettersen, E.F. et al. UCSF ChimeraX: Structure visualization for researchers, educators, and developers. Protein Sci 30, 70–82 (2021).

57. Schrodinger, L., DeLano, W. PyMOL [Internet]. Available at: http://www.pymol.org/pymol. (2020).

58. Marapana, D.S. et al. Plasmepsin V cleaves malaria effector proteins in a distinct endoplasmic reticulum translocation interactome for export to the erythrocyte. Nat. Microbiol. 3, 1010–1022 (2018).

59. Marapana, D.S. et al. GID/CTLH E3 ligase complex control cell fate programs for sexual development of *Plasmodium falciparum*. Nat Commun 17 (2026).

60. Bianco, A.E. et al. A repetitive antigen of *Plasmodium falciparum* that is homologous to heat shock protein 70 of *Drosophila melanogaste*r. Proc. Natl. Acad. Sci. USA 83, 8713–8717 (1986).

