## Extended Data Figures and legends for "Molecular basis for the role of Ripr in *Plasmodium falciparum* invasion of human erythrocytes"

Running title: EGF7-mediated PCRCR disruption of *P. falciparum* invasion

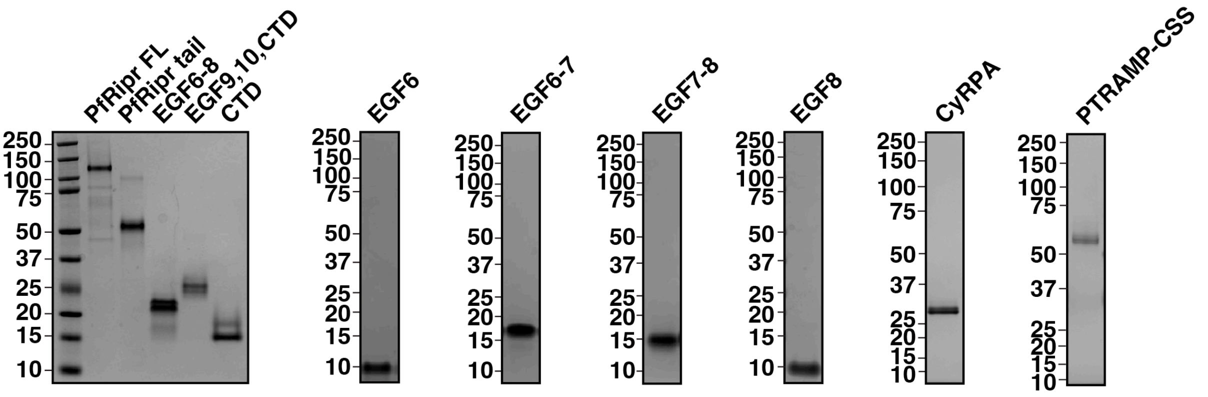


**Extended Data Fig. 1.** Non-reduced Coomassie stained SDS-PAGE of PfRipr domains expressed as recombinant proteins. From left to right: molecular weight markers; full-length PfRipr (PfRipr FL) [1]; PfRipr tail; PfRipr$^{EGF6-8}$ (EGF6-8); PfRipr$^{EGF9,10,CTD}$ (EGF9,10,CTD); PfRipr$^{CTD}$(CTD); PfRipr$^{EGF6}$ (EGF6); PfRipr$^{EGF6-7}$ (EGF6-7); PfRipr$^{EGF7-8}$ (EGF7-8); PfRipr$^{EGF8}$ (EGF8). Also shown are recombinant PfCyRPA and PfPTRAMP-CSS [2].

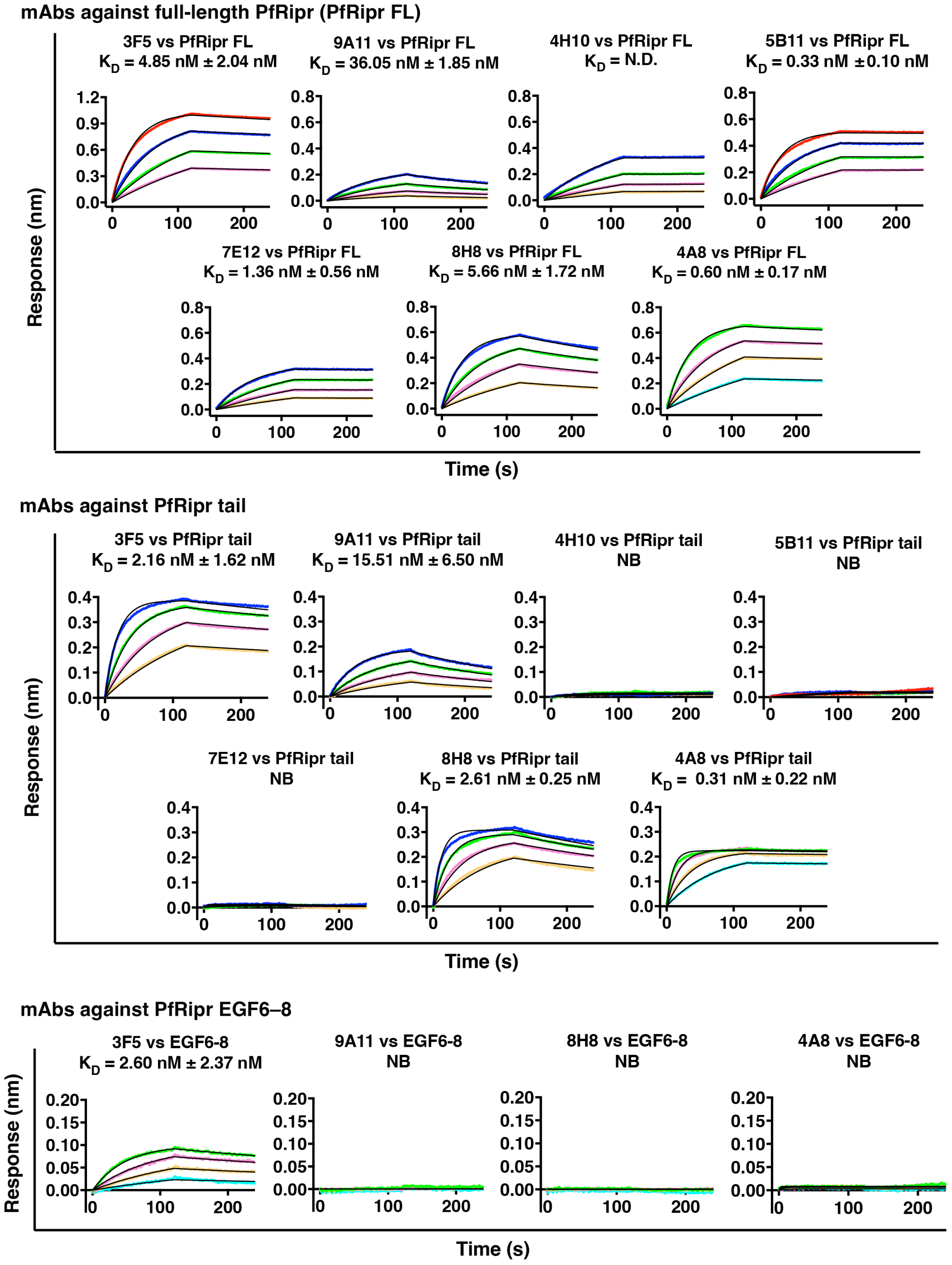


**Extended Data Fig. 2.** Kinetics studies and binding affinities of PfRipr-specific mAbs to full-length PfRipr (PfRipr FL) and PfRipr$^{tail}$; and PfRipr$^{EGF6–8}$ for PfRipr$^{tail}$-specific mAbs measured using BLI. Antibodies are immobilised onto anti-mouse Fc (AMC) biosensors and associated with serially diluted PfRipr antigens, and dissociation rate monitored when biosensors were replaced in kinetics buffer. Concentrations of antigens are colour coded: 500 nM (orange), 250 nM (red), 125 nM (dark blue), 62.5 nM (green), 31.25 nM (pink), 15.63 nM (yellow), 7.81 nM (light blue). Representative sensorgrams (association and dissociation) and 1:1 model best fit (black) is representative of two independent measurements. Reported K$_D$ indicates mean ± SEM. NB: No binding. N.D: not determinable due to slow dissociation rate.

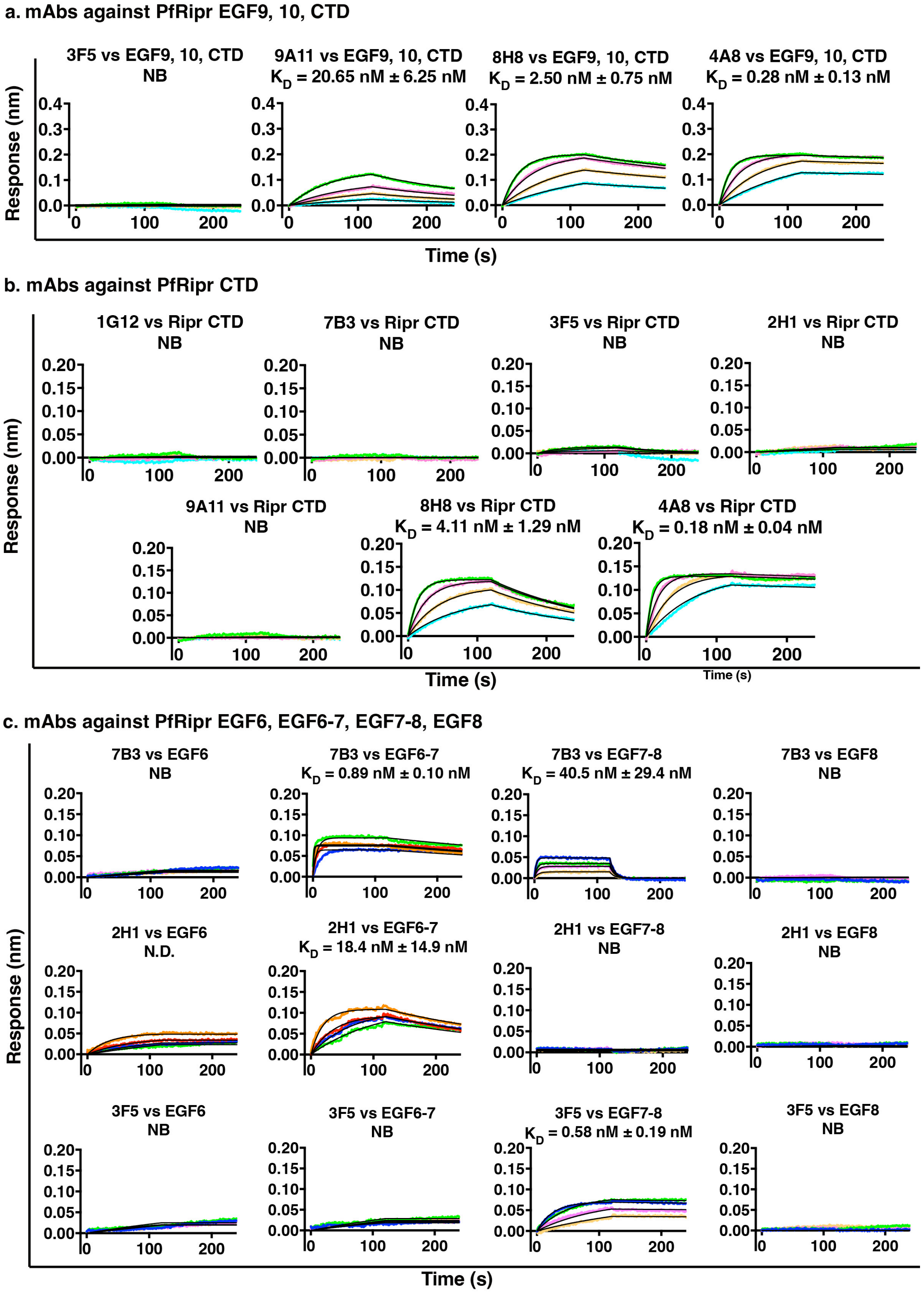


**Extended Data Fig. 3.** Kinetics studies and binding affinities of PfRipr$^{tail}$-specific mAbs to PfRipr$^{9\text{-}10\text{-}CTD}$ and PfRipr$^{CTD}$; and PfRipr$^{EGF6}$, PfRipr$^{EGF6–7}$, PfRipr$^{EGF7–8}$, PfRipr$^{EGF8}$ for PfRipr$^{EGF6–8}$-specific mAbs, measured using BLI, following the same protocol and colour-coding as in **Extended Data Fig. 2.**

| mAb2 / mAb1 | CyRPA | 5B11 | 6D2 | 7E12 | 4H10 | 1G12 | 7B3 | 5B3 | 3F5 | 8H8 | 2H1 | 9A11 | PTRAMP CSS | 4A8 |
|---|---|---|---|---|---|---|---|---|---|---|---|---|---|---|
| 5B11 | 25 | 14 | 101 | 99 | 72 | 98 | 96 | 95 | 98 | 100 | 105 | 90 | 151 | 176 |
| 6D2 | 132 | 89 | 18 | 62 | 96 | 102 | 95 | 93 | 94 | 99 | 108 | 93 | 155 | 176 |
| 7E12 | 103 | 81 | 52 | 15 | 64 | 106 | 90 | 90 | 94 | 85 | 72 | 69 | 64 | 89 |
| 4H10 | 131 | 73 | 44 | 53 | 24 | 76 | 56 | 79 | 78 | 65 | 52 | 42 | 159 | 77 |
| 1G12 | 136 | 97 | 98 | 114 | 96 | 17 | 28 | 61 | 97 | 99 | 93 | 100 | 117 | 215 |
| 7B3 | 133 | 84 | 85 | 78 | 82 | -13 | 19 | 40 | 62 | 63 | 100 | 67 | 95 | 170 |
| 3F5 | 135 | 70 | 70 | 82 | 59 | 100 | 78 | 2 | 12 | 83 | 65 | 67 | 187 | 84 |
| 8H8 | 130 | 52 | 53 | 52 | 53 | 96 | 82 | 72 | 84 | 14 | 57 | 51 | 68 | 78 |
| 2H1 | 129 | 77 | 75 | 60 | 68 | 65 | 93 | 76 | 63 | 64 | 12 | 29 | 76 | 131 |
| 9A11 | 118 | 75 | 70 | 59 | 53 | 103 | 86 | 87 | 78 | 63 | 67 | 18 | -101 | 83 |
| PTRAMP CSS | | 83 | 84 | 94 | 93 | 81 | 82 | 82 | 96 | 95 | 83 | 77 | | 88 |
| 4A8 | 127 | | | | | | | 70 | | | | | 34 | 31 |

**Extended Data Fig. 4. Competition studies of anti-PfRipr monoclonal antibodies.** Epitope binning matrix of anti-PfRipr mAbs and competition assay with PfPTRAMP-CSS and PfCyRPA. PfRipr with C-terminal His-tag was immobilised onto Ni-NTA biosensors, followed by association with anti-PfRipr Abs or PfPTRAMP-CSS (mAb1 indicated on the left column), and binding to secondary mAb or PfPTRAMP-CSS or PfCyRPA (mAb2 shown on top row). Reported competition scores are obtained by dividing the individual mAb2 responses by the kinetics buffer control signal, multiplied by 100. A score below 34 indicates competition; 34 and above indicates no competition. Scores that indicate competition are coloured in red. Recombinant PfPTRAMP-CSS used was produced as previously described [2]. PfCyRPA competition assay was carried out one way with PfCyRPA-Rh5 as the secondary Ag. Epitope binning and competition assays for mAb 5B3 were also carried out one way as the secondary mAb. 4A8 as primary mAb displayed a fast off-rate, resulting in intermediate binding values for true non-competing antibodies. Therefore, these data were excluded from the analysis (grey), except when 5B3, PfCyRPA and PfPTRAMP-CSS were used as a secondary.

**a**

| mAbs | $K_D$ (nM) | $k_a$ (1/Ms) | $k_d$ (1/s) |
|---|---|---|---|
| 1G12 | 1.02 ± 0.40 | $1.96\times10^{5}\pm 8.15\times10^{4}$ | $1.68\times10^{-4}\pm 5.9\times10^{-6}$ |
| 7B3 | 3.43 ± 1.65 | $2.36\times10^{5}\pm 8.26\times10^{4}$ | $6.73\times10^{-4}\pm 1.07\times10^{-4}$ |
| 2H1 | 15.31± 7.20 | $1.90\times10^{5}\pm 6.33\times10^{4}$ | $2.45\times10^{-3}\pm 3.94\times10^{-4}$ |
| 3F5 | 4.85 ± 2.04 | $1.02\times10^{5}\pm 3.88\times10^{4}$ | $4.14\times10^{-4}\pm 1.94\times10^{-5}$ |
| 5B11 | 0.33 ± 0.10 | $2.27\times10^{5}\pm 9.09\times10^{4}$ | $6.50\times10^{-5}\pm 6.98\times10^{-6}$ |
| 6D2 | 1.09 ± 0.72 | $6.24\times10^{5}\pm 2.93\times10^{4}$ | $4.73\times10^{-5}\pm 1.29\times10^{-5}$ |
| 7E12 | 1.36 ± 0.56 | $1.01\times10^{5}\pm 2.61\times10^{4}$ | $1.23\times10^{-4}\pm 2.08\times10^{-5}$ |
| 8H8 | 5.66 ± 1.72 | $3.51\times10^{5}\pm 1.09\times10^{5}$ | $1.80\times10^{-3}\pm 1.05\times10^{-5}$ |
| 9A11 | 36.05 ± 1.85 | $9.01\times10^{5}\pm 4.93\times10^{3}$ | $3.26\times10^{-3}\pm 3.44\times10^{-4}$ |
| 4H10 | N.D | $1.03\times10^{5}\pm 1.59\times10^{4}$ | N.D |
| 4A8 | 0.60 ± 0.17 | $6.73\times10^{5}\pm 1.95\times10^{5}$ | $3.71\times10^{-4}\pm 4.15\times10^{-6}$ |
| 5B3* | 0.59 | $2.29\times10^{5}$ | $1.34\times10^{-4}$ |

**b**

| mAbs | $K_D$ (nM) | $k_a$ (1/Ms) | $k_d$ (1/s) |
|---|---|---|---|
| 1G12 | 0.98 ± 0.80 | $3.22\times10^{5}\pm 5.50\times10^{2}$ | $3.16\times10^{-4}\pm 2.56\times10^{-4}$ |
| 7B3 | 2.85 ± 0.20 | $3.59\times10^{5}\pm 2.00\times10^{4}$ | $1.02\times10^{-3}\pm 1.40\times10^{-5}$ |
| 2H1 | 7.73 ± 1.41 | $4.93\times10^{5}\pm 1.00\times10^{5}$ | $3.27\times10^{-3}\pm 3.16\times10^{-4}$ |
| 3F5 | 2.16 ± 1.62 | $3.82\times10^{5}\pm 1.68\times10^{5}$ | $5.52\times10^{-4}\pm 2.59\times10^{-4}$ |
| 8H8 | 2.61 ± 0.25 | $7.95\times10^{5}\pm 1.32\times10^{5}$ | $2.04\times10^{-3}\pm 1.48\times10^{-4}$ |
| 9A11 | 15.51 ± 6.50 | $2.11\times10^{5}\pm 2.59\times10^{4}$ | $3.10\times10^{-3}\pm 9.67\times10^{-4}$ |
| 4A8 | 0.31 ± 0.22 | $1.49\times10^{5}\pm 2.68\times10^{5}$ | $4.06\times10^{-4}\pm 2.46\times10^{-4}$ |

**c**

| | EGF6-8 | | | EGF9-10,CTD | | | CTD | | |
|---|---|---|---|---|---|---|---|---|---|
| | $K_D$ (nM) | $k_a$ (1/Ms) | $k_d$ (1/s) | $K_D$ (nM) | $k_a$ (1/Ms) | $k_d$ (1/s) | $K_D$ (nM) | $k_a$ (1/Ms) | $k_d$ (1/s) |
| 1G12 | 2.29 ± 1.8 | $2.63\times10^{5}$ | $6.31\times10^{-4}$ | | | | | | |
| 7B3 | 2.93 ± 0.39 | $2.88\times10^{5}$ | $9.58\times10^{-4}$ | | | | | | |
| 2H1 | 10.84 ± 5.86 | $5.93\times10^{5}$ | $2.95\times10^{-3}$ | | | | | | |
| 3F5 | 2.60 ± 2.37 | $3.08\times10^{5}$ | $1.53\times10^{-3}$ | | | | | | |
| 9A11 | | | | 20.65 ± 6.25 | $1.91\times10^{5}$ | $5.12\times10^{-3}$ | | | |
| 8H8 | | | | 2.50 ± 0.75 | $6.31\times10^{5}$ | $2.05\times10^{-3}$ | 4.11 ± 1.29 | $1.06\times10^{6}$ | $5.73\times10^{-3}$ |
| 4A8 | | | | 0.28 ± 0.13 | $1.06\times10^{6}$ | $4.39\times10^{-4}$ | 0.18 ± 0.04 | $1.84\times10^{6}$ | $3.95\times10^{-4}$ |

**Extended Data Fig. 5. Binding kinetics of anti-PfRipr mAbs. a.** Binding kinetics of anti-PfRipr mAbs to full-length PfRipr. N = 2. Parameters shown as mean ± SEM. N.D. non-determinable due to slow dissociation-rate. Kinetics parameters for 5B3 are shown against full-length PkRipr. N = 1. **b.** Binding kinetics of anti-PfRipr$^{tail}$ mAbs to PfRipr$^{tail}$. N = 2. Parameters shown as mean ± SEM. **c.** Binding kinetics of anti-PfRipr$^{tail}$ mAbs to PfRipr tail constructs. N = 2. $K_D$ data shown as mean ± SEM. $k_a$ and $k_d$ data shown as mean of the duplicates.

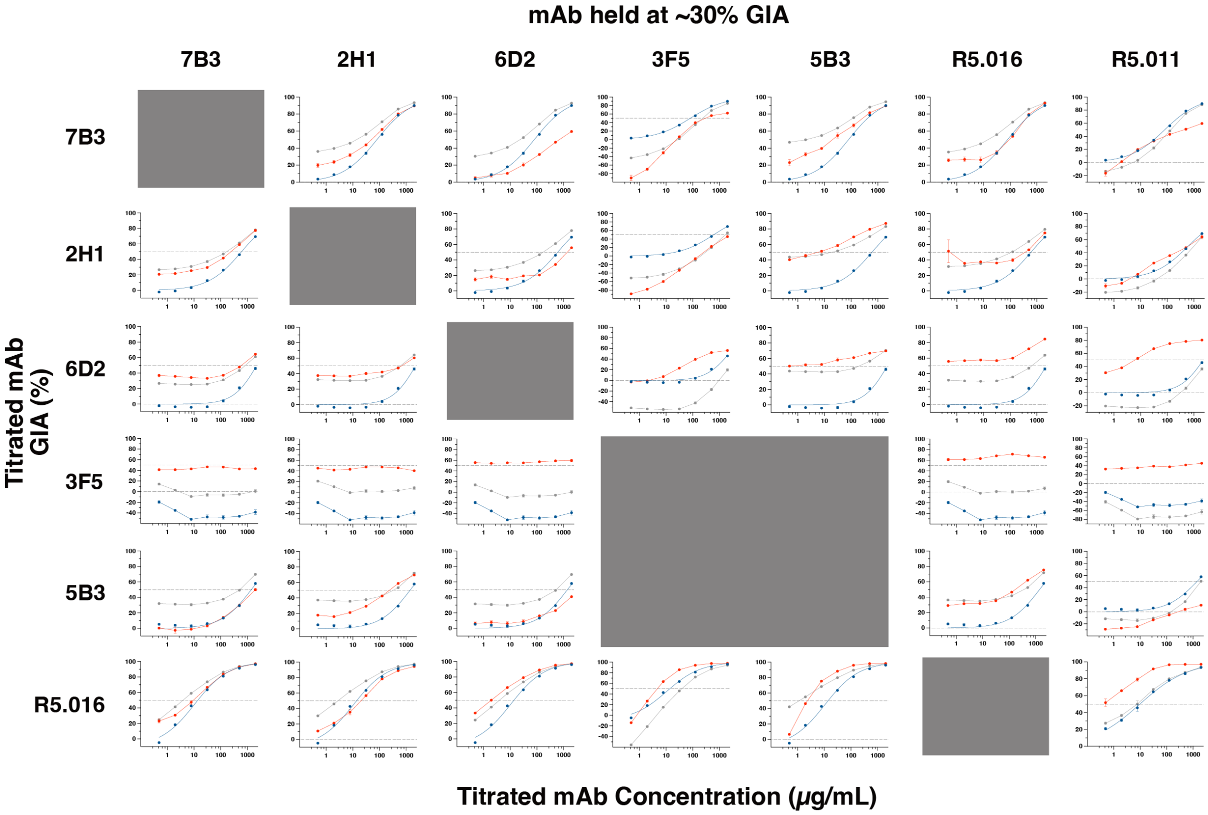


**Extended Data Fig. 6. Complete pairwise synergy titration matrix for anti-PfRipr and anti-Rh5 mAb combinations.** Full matrix of pairwise combinatorial GIA titration curves. Rows indicate the titrated mAb (four-fold seven-step serial dilution from 2 mg/mL); columns indicate the mAb held constant at approximately 30% GIA (or 500 μg/mL for 3F5 and R5.011). Predicted Bliss additivity (grey) is compared with measured GIA (red) for each combination. Blue curves represent titrated mAb alone. Grey-shaded cells indicate combinations where the held mAb competes with the titrated mAb and were not tested. Individual points are the mean of a triplicate measurement with error bars indicating SEM. Source data are provided as a Source Data file.

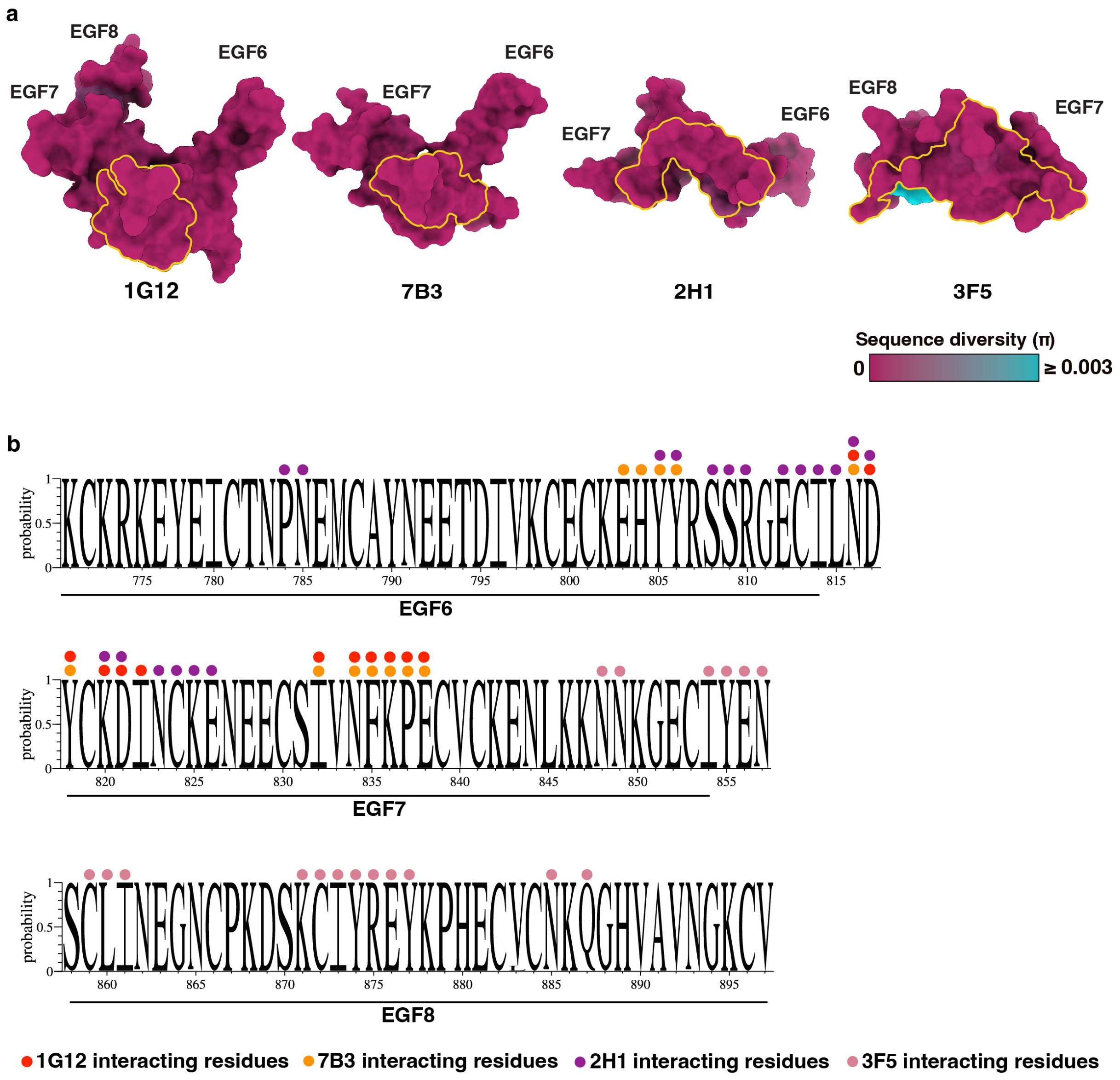


**Extended Data Fig. 7. Sequence conservation of Ripr$^{EGF6–8}$ across *P. falciparum* strains mapped onto the antigen of the Fab-EGF complexes. a.** Sequence conservation of Ripr$^{EGF6-8}$ across *P. falciparum* strains mapped onto the antigen of the Fab-EGF complexes. Surface representation of PfRipr$^{EGF6-8}$ coloured according to sequence diversity (π) among *P. falciparum* strains sequenced, ranging from 0 (fully conserved, magenta) to ≥ 0.003 (highly polymorphic, cyan). Antibody epitopes are outlined in yellow. From left to right: 1G12-EGF6-8, 7B3-EGF6-7, 2H1-EGF6-7 and 3F5-EGF7-8 (from 20,646 *P. falciparum* strains). **b.** Sequence diversity of PfRipr$^{EGF6-8}$ subdomains across 20,646 *P.falciparum* strains sequenced in Weblogo representation (*38*). 1G12, 7B3, 2H1 and 3F5 interacting residues are indicated in circles coloured according to Fig. 3a.

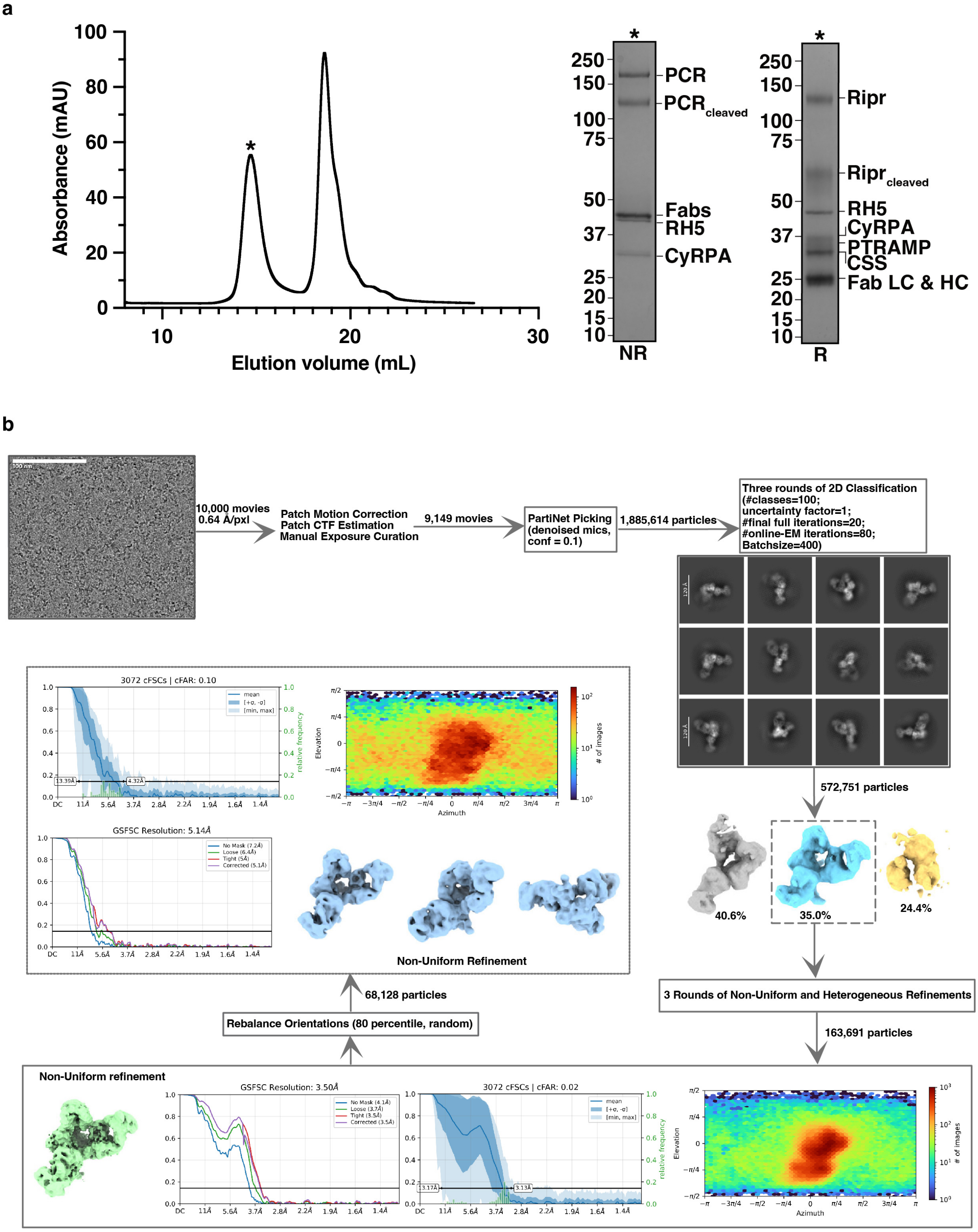


**Extended Data Fig. 8. a.** Size-exclusion chromatography traces (Superose 6 Increase GL 10/300) of the stabilised PfPCRCR+7B3+2H1+3F5+6D2 Fabs co-complex. Peak fraction (as indicated by asterisk) was analyzed by non-reducing (NR) and reducing (R) SDS-PAGE confirming co-elution of all components. PCR stands for the stabilised PfPTRAMP-CSS-Ripr co-complex. LC and HC stand for the light chain and heavy chain of Fab fragments respectively. **b.** Cryo-EM image processing workflow for the PfPCRCR-Fabs co-complex. A total of 10,000 movies collected at 0.64 Å/pixel were processed in CryoSPARC. Following manual exposure curation, 9,149 micrographs were retained for downstream processing. PartiNet particle picking on denoised micrographs yielded 1,885,614 particles, which were subjected to three rounds of reference-free 2D classification, resulting in 572,751 particles for ab initio reconstruction into three classes. The best-resolved class was refined through iterative heterogeneous and non-uniform refinement. To alleviate severe preferred orientation observed in the initial 3.50 Å reconstruction, orientation rebalancing was applied, resulting in a final subset of 68,128 particles used for the final 5.14 Å reconstruction (EMDB-80770). Representative 2D class averages, density maps, directional FSC plots, and angular distribution plots are shown.

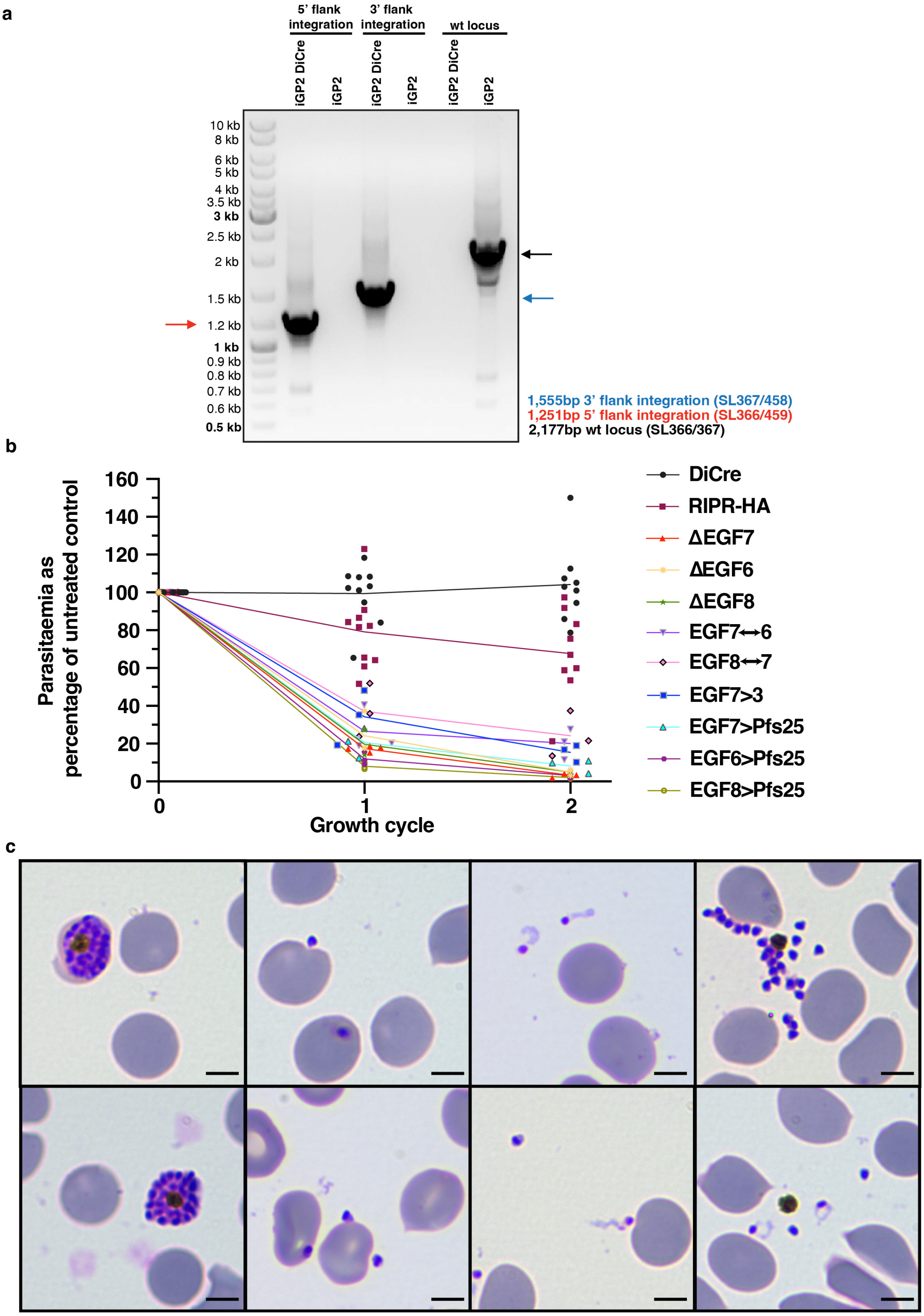


**Extended Data Fig. 9. a.** PCR validation of iGP2-DiCre cell line creation. Blue, red and black arrowheads indicate the expected 3', 5' and wild type PCR-amplified gene products respectively. **b.** Growth assays of all transgenic parasites over one and two replication cycles (x-axis), shown as parasitaemia normalised to untreated (– rapamycin) controls of the same line at the same cycle (y-axis). Except RIPR–HA line, any remaining parasites in rapamycin-treated transgenic lines at cycle 2 were committed gametocytes carried over from the starting culture; no asexual-stage parasites were detected. Data presented using Prism v10 (GraphPad). **c.** Representative Giemsa images of ΔEGF7 transgenic parasites at the end of the same intraerythrocytic cycle following rapamycin treatment. Scale bar = 4μm.

### References

1. Wong, W. *et al.* Structure of *Plasmodium falciparum* Rh5-CyRPA-Ripr invasion complex. *Nature* **565**, 118-121 (2019).
2. Seager, B.A. *et al.* PTRAMP, CSS and Ripr form a conserved complex required for merozoite invasion of Plasmodium species into erythrocytes. *Nat Commun* **17**, 1780 (2026).
