## Supplementary material for "Molecular basis for the role of Ripr in *Plasmodium falciparum* invasion of human erythrocytes"

Running title: EGF7-mediated PCRCR disruption of *P. falciparum* invasion

### Supplementary Material

#### Table S1. Anti-PfRipr antibody gene features

| mAb | IGHV | IGHD | IGHJ | HCDR3 | IGLV | IGLJ | LCDR3 |
|---|---|---|---|---|---|---|---|
| **1G12*** | IGHV1-71-11*01 | IGHD1-2*01 | IGHJ3*01 | ARNYFGYEGFAY | IGKV1-117*01 | IGKJ2*01 | FQHSHFPYT |
| **2H1** | IGHV2-9*02 | IGHD2*01 | IGHJ4*01 | VRDYYGRYYYAMDN | IGLV7-46*01 | IGLJ3*02 | ALWYSNHWV |
| **3F5** | IGHV9-3-1*01 | IGHD3-1*01 | IGHJ4*01 | ARPSSGYGYYSLDY | IGKV15-103*01 | IGKJ5*01 | QQGQSYPFT |
| **4H10** | IGHV5-6*01 | IGHD2-2*01 | IGHJ4*01 | VRLVPTGHHAMDY | IGKV19-93*01 | IGKJ1*01 | LQYDNLLRT |
| **5B3**[#] | IGHV14-1*02 | IGHD1-1*01 | IGHJ4*01 | ARSYYYGSSDAMDN | IGKV4-72*01 | IGKJ1*01 | QQWSPNPWT |
| **5B11** | IGHV1-47-36*01 | IGHD1-1*01 | IGHJ1*01 | ARSRFYGTSPYWYFDV | IGKV14-111*01 | IGKJ2*01 | LQYDEFPYT |
| **6D2** | IGHV1-47-11*01 | N/A | IGHJ2*01 | ARGGNYFDY | IGKV4-68*01 | IGKJ5*01 | QLWTSNPLT |
| **7B3** | IGHV1-42-1*01 | IGHD1-2*01 | IGHJ3*01 | APDYGYVGFAY | IGKV1-117*01 | IGKJ1*01 | FQGSHVPRT |
| **7E12** | IGHV1-69-2*01 | N/A | IGHJ4*01 | ARDAMDY | IGLV1*01 | IGLJ*01 | ALWYSNHWV |
| **8H8** | IGHV1-47-46*01 | IGHD1-1*01 | IGHJ1*01 | TRSGYYGTYWYFDV | IGKV3-5*01 | IGKJ*01 | QQSNEDPPT |
| **9A11** | IGHV1S135*01 | IGHD1-1*02 | IGHJ2*01 | ARGGTNFDY | IGKV3-2*01 | IGKJ1*01 | QQSKEVPWT |

*Healer et al., 2019 [1]
[#]Seager et al., 2026 [2]

### Table S2. Data collection and refinement statistics

| | 1G12-EGF6–8 | 7B3-EGF6–8 | 2H1-EGF6–7 | 3F5-EGF7–8 |
|---|---|---|---|---|
| **Beamline** | MX2 | MX2 | MX2 | MX2 |
| **Wavelength (Å)** | 0.95365 | 0.95365 | 0.95365 | 0.95365 |
| **Space group** | $C222_1$ | I222 | $P22_12_1$ | $P2_12_12_1$ |
| **Cell dimensions** | | | | |
| ***a,b,c* (Å)** | 74.8, 277.5, 90.6 | 73.7, 90.2 252.7 | 74.0, 117.0, 134.8 | 98.7, 120.1, 219.6 |
| **α, β, γ (°)** | 90, 90, 90 | 90, 90, 90 | 90, 90, 90 | 90, 90, 90 |
| **Resolution (Å)[a]** | 39.99-2.80 (2.95-2.80) | 45.11-2.99 (3.11-2.99) | 45.89-2.69 (2.82-2.69) | 48.16-3.43 (3.59-3.43) |
| **No. molecules in ASU** | 1 | 1 | 2 | 4 |
| **No. observations** | 131,283 (17,925) | 64,685 (3496) | 186,826 (23,790) | 242,714 (33,303) |
| **No. unique observations** | 23,732 (3,402) | 13,157 (658) | 33,257 (4,356) | 35,906 (4,697) |
| **Multiplicity** | 5.5 (5.3) | 4.9 (5.3) | 5.6 (5.5) | 6.8 (7.1) |
| **$R_{merge}$ (%)** | 19.7 (105.3) | 20.1 (83.0) | 16.3 (98.1) | 24.6 (112.5) |
| **$R_{pim}$ (%)** | 9.9 (54.3) | 10.0 (40.5) | 8.3 (49.9) | 10.9 (48.8) |
| **$\langle I/\sigma I \rangle$** | 7.2 (1.7) | 7.7 (2.2) | 9.8 (1.8) | 5.7 (1.9) |
| **$CC_{½}$** | 98.8 (63.6) | 99.0 (72.4) | 99.1 (51.6) | 99.4 (79.4) |
| **Completeness (%)** | 99.9 (100.0) | 93.4 (85.1) | 100.0 (100.0) | 99.9 (99.9) |
| **Refinement Statistics** | | | | |
| **Reflections (work)** | 23,700 | 13,091 | 33,178 | 35,673 |
| **Reflections (test)** | 1,198 | 1,310 | 1,625 | 1,810 |
| **Non-hydrogen atoms** | 4,317 | 3,982 | 7,931 | 14,974 |
| **Macromolecule** | 4,192 | 3,938 | 7,676 | 14,966 |
| **Solvent** | 51 | 20 | 175 | 0 |
| **Heteroatom** | 74 | 24 | 80 | 8 |
| **$R_{work}$ / $R_{free}$[e]** | 19.9 / 23.5 | 21.2 / 26.8 | 18.5 / 24.2 | 24.0 / 28.5 |
| **Rms deviations from ideality** | | | | |
| **Bond lengths (Å)** | 0.002 | 0.002 | 0.004 | 0.002 |
| **Bond angle (°)** | 0.49 | 0.46 | 0.73 | 0.57 |
| **Ramachandran plot** | | | | |
| **Favoured regions (%)** | 93.8 | 92.1 | 96.1 | 95.2 |
| **Allowed regions (%)** | 6.2 | 7.9 | 3.9 | 4.8 |
| **B-factors ($Å^2$)** | | | | |
| **Wilson B-value** | 50.5 | 46.0 | 39.5 | 75.6 |
| **Average B-factors** | 62.9 | 55.7 | 50.6 | 91.6 |
| **Average macromolecule** | 63.1 | 55.9 | 50.7 | 91.6 |
| **Average heteroatom** | 65.2 | 50.5 | 66.2 | 98.2 |
| **Average solvent** | 43.5 | 19.4 | 40.6 | - |

[a] Values in parentheses refer to the highest resolution bin.

[e] 5% of data were used for the $R_{free}$ calculation

**Table S3. Table of contents between 1G12 and PfRipr EGF6-8.**

| EGF6-8 Residue (BSA Å$^2$) | Interaction Type | 1G12 Residue |
|---|---|---|
| **Asn816 (6.4)** | | |
| Asn | VDW | H-Tyr96 |
| **Asp817 (39.3)** | | |
| Asp | VDW | H-Tyr96, H-Tyr99 |
| Asp$^{N}$ | HB | H-Tyr96$^{OH}$ |
| **Tyr818 (63.77)** | | |
| Tyr | VDW | H-Tyr96, H-Phe97, H-Tyr99 |
| Tyr$^{N}$ | HB | H-Tyr96$^{OH}$ |
| **Lys820 (17.49)** | | |
| Lys | VDW | H-Tyr99 |
| **Asp821 (115.48)** | | |
| Asp | VDW | K-Asn28, K-Asn30, K-Tyr32, K-Lys50 H-Phe97, H-Gly98, H-Tyr99 |
| Asp$^{OD1}$ | SB, HB | K-Lys50$^{NZ}$ |
| Asp$^{OD2}$ | SB | K-Lys50$^{NZ}$ |
| Asp$^{O}$ | HB | K-Asn30$^{ND2}$ |
| **Ile822 (33.89)** | | |
| Ile | VDW | K-Asn28, K-Asn30 |
| **Ile832 (26.97)** | | |
| Ile | VDW | H-Phe97 |
| **Asn834 (11.34)** | | |
| Asn | VDW | H-Asn58 |
| **Asn834 (28.10)** | | |
| Asn | VDW | K-Phe94, K-Tyr96 |
| **Phe835 (158.87)** | | |
| Phe | VDW | H-Leu33, H-Phe97, H-Val50, H-Ile51, H-Asn52, H-Asp56, H-Thr57, H-Asn58 |
| **Phe835 (25.30)** | | |
| Phe | VDW | K-Phe94, K-Tyr96 |
| **Lys836 (31.33)** | | |
| Lys | VDW | H-Phe97, H-Gly98, H-Asn95 |
| **Lys836 (103.78)** | | |
| Lys | VDW | K-His27D, K-Asn28, K-Tyr32, K-His91, K-Tyr96 |
| Lys$^{NZ}$ | HB | K-His91$^{O}$, K-Tyr96$^{OH}$ |
| **Pro837 (9.40)** | | |
| Pro | VDW | H-Phe97 |
| **Pro837 (23.80)** | | |
| Pro | VDW | K-His27D, K-Asn28, K-Tyr32 |
| Pro$^{O}$ | HB | K-Asn28$^{ND2}$ |
| **Glu838 (27.12)** | | |
| Glu | VDW | K-His27D, K-Ser27E, K-Asn28 |

**Table S4. Table of contents between 7B3 and PfRipr EGF6-8.**

| EGF6-8 Residue (BSA Å$^2$) | Interaction Type | 7B3 Residue |
|---|---|---|
| **Glu803 (66.16)** | | |
| Glu | VDW | H-Tyr53, H-Arg52, H-Asn54 |
| Glu$^{OE2}$ | SB | H-Arg52$^{NH1}$ |
| Glu$^{OE2}$ | SB, HB | H-Arg52$^{NH2}$ |
| **His804 (3.30)** | | |
| His | VDW | H-Arg52, H-Asp31 |
| **Tyr805 (7.37)** | | |
| Tyr | VDW | H-Asp31 |
| **Tyr806 (9.18)** | | |
| Tyr | VDW | H-Asp31 |
| **Asn816 (22.56)** | | |
| Asn | VDW | H-Asp31 |
| Asn$^{ND2}$ | HB | H-Asp31$^{OD2}$ |
| **Tyr818 (64.88)** | | |
| Tyr | VDW | H-Tyr32, H-Asp31, H-Gly97, H-Tyr98 |
| Tyr$^{OH}$ | HB | H-Asp31$^{OD1}$ |
| **Ile832 (56.97)** | | |
| Ile | VDW | H-Arg52, H-Tyr98 |
| **Val833 (11.76)** | | |
| Val | VDW | K-His27D, K-Asn28 |
| Val$^{O}$ | HB | K-His27D$^{NE2}$ |
| **Asn834 (47.54)** | | |
| Asn | VDW | K-His27D, K-Ser27E, K-Arg96 |
| **Phe835 (176.85)** | | |
| Phe | VDW | K-Val94, K-Arg96<br>H-Thr33, H-Tyr98, H-Arg52, H-Asp56, H-Thr57, H-Gly58, H-Leu50, H-Asp95 |
| **Lys836 (138.08)** | | |
| Lys | VDW | K-His27D, K-Asn28, K-Tyr32, K-Arg96, K-Gly91<br>H-Tyr98, H-Asp95, H-Val99 |
| Lys$^{NZ}$ | SB | H-Asp95$^{OD1}$ |
| Lys$^{NZ}$ | SB, HB | H-Asp95$^{OD2}$ |
| Lys$^{NZ}$ | HB | H-Tyr98$^{O}$ |
| **Pro837 (45.35)** | | |
| Pro | VDW | K-Asn28, K-Tyr32<br>H-Tyr98 |
| Pro$^{O}$ | HB | K-Asn28$^{ND2}$ |
| **Glu838 (49.53)** | | |
| Glu | VDW | K-Ser27E, K-Asn28 |
| Glu$^{OE2}$ | HB | K-Asn28$^{N}$ |

**Table S5. Table of contacts between 2H1 and PfRipr EGF6-7.**

| EGF6-7 Residue (BSA Å$^2$) | Interaction Type | 2H1 Residue |
|---|---|---|
| **Pro784 (43.52)** | | |
| Pro | VDW | H-Thr30, H-Asn73, |
| **Asn785 (28.97)** | | |
| Asn | VDW | H-Thr30, H-Asn31 |
| Asn$^{ND2}$ | HB | H-Thr30$^{O}$, H-Thr30$^{OG1}$, H-Asn31$^{OD1}$ |
| **Tyr805 (3.96)** | | |
| Tyr | VDW | H-Asn31 |
| **Tyr806 (31.66)** | | |
| Tyr | VDW | H-Tyr97, H-Gly98, H-Arg99 |
| Tyr$^{OH}$ | HB | H-Tyr97$^{O}$, H-Gly98$^{O}$, H-Arg99$^{N}$ |
| **Ser808 (9.09)** | | |
| Ser | VDW | H-Tyr97 |
| Ser$^{OG}$ | HB | H-Tyr97$^{OH}$ |
| **Ser809 (6.01)** | | |
| Ser | VDW | H-Tyr97, H-Tyr100B |
| **Arg810 (140.38)** | | |
| Arg | VDW | L-Trp91<br>H-Tyr97, H-Trp52, H-Ser56,<br>H-Tyr100B, H-Asp58, |
| Arg$^{NE}$ | HB | H-Tyr97$^{OH}$, |
| Arg$^{NH1}$ | HB, SB | H-Asp58$^{OD2}$ |
| Arg$^{NH1}$ | HB | H-Trp52$^{NE1}$ |
| Arg$^{NH2}$ | HB | H-Tyr100B$^{OH}$ |
| **Glu812 (61.25)** | | |
| Glu | VDW | H-Trp52, H-Pro53, H-Gly54, H-Gly55, H-Tyr97 |
| Glu$^{OE1}$ | HB | H-Gly54$^{N}$ |
| Glu$^{OE2}$ | HB | H-Tyr97$^{OH}$ |
| **Cys813 (0.98)** | | |
| Cys | VDW | H-Thr30, H-Asn31 |
| **Ile814 (46.40)** | | |
| Ile | VDW | H-Thr30, H-Asn31, H-Tyr97, H-Pro53 |
| **Leu815 (22.32)** | | |
| Leu | VDW | H-Asn31 |
| Leu$^{N}$ | HB | H-Asn31$^{OD1}$ |
| **Asn816 (8.71)** | | |
| Asn | VDW | H-Arg99 |
| **Asp817 (28.91)** | | |
| Asp | VDW | H-Tyr96, H-Tyr97, H-Arg99, H-Gly98, H-Tyr100, H-Tyr100A |
| Asp$^{O}$ | HB | H-Tyr96$^{OH}$ |
| Asp$^{OD1}$ | HB | H-Arg99$^{N}$ |
| Asp$^{OD2}$ | HB | H-Arg99$^{N}$ |

| | | |
|---|---|---|
| **Lys820 (52.65)** | | |
| Lys | VDW | H-Asn31, H-Tyr32, H-Tyr96, H-Tyr97, H-Gly98 |
| $Lys^{NZ}$ | HB | H-Asn31$^{O}$, H-Tyr96$^{O}$, H-Tyr97$^{O}$ |
| **Asp821 (17.08)** | | |
| Asp | VDW | H-Tyr96 |
| $Asp^{OD1}$ | HB | H-Tyr96$^{OH}$ |
| **Asn823 (31.48)** | | |
| Asn | VDW | H-Gly26, H-Val2, H-Arg94 |
| **Cys824 (13.49)** | | |
| Cys | VDW | H-Gly26 |
| **Lys825 (43.01)** | | |
| Lys | VDW | H-Gln1 |
| **Glu826 (42.68)** | | |
| Glu | VDW | H-Ser25, H-Gly26, H-Gln1 |
| $Glu^{OE2}$ | HB | H-Gly26$^{N}$ |

**Table S6. Table of contents between 3F5 and PfRipr EGF7-8.**

| EGF7-8 Residue (BSA Å$^2$) | Interaction Type | 3F5 Residue |
|---|---|---|
| **Asn848 (41.13)** | | |
| Asn | VDW | H-Thr28, H-Tyr27, H-Gly26 |
| Asn$^{ND2}$ | HB | H-Thr28$^{OG1}$ |
| **Asn849 (15.89)** | | |
| Asn | VDW | H-Gly26 |
| **Ile854 (19.07)** | | |
| Ile | VDW | H-Thr28, H-Asn31 |
| **Tyr855 (0.98)** | | |
| Tyr | VDW | H-Asn31 |
| Tyr$^{O}$ | HB | H-Asn31$^{ND2}$ |
| **Glu856 (51.41)** | | |
| Glu | VDW | H-Thr30, H-Asn31, H-Tyr53 |
| Glu$^{OE2}$ | HB | H-Asn31$^{OD1}$ |
| **Asn857 (40.14)** | | |
| Asn | VDW | H-Thr30, H-Asn31, H-Tyr32, H-Gly33, H-Tyr100A |
| Asn$^{ND2}$ | HB | H-Asn31$^{O}$ |
| **Cys859 (15.90)** | | |
| Cys | VDW | H-Tyr100A |
| **Leu860 (138.60)** | | |
| Leu | VDW | H-Thr30, H-Asn31, H-Tyr32, H-Gly33, H-Trp50, H-Asn52, H-Tyr53, H-Thr54, H-Tyr100A |
| Leu$^{O}$ | HB | H-Asn52$^{ND2}$ |
| **Ile861 (42.00)** | | |
| Ile | VDW | H-Tyr53 |
| **Lys871 (128.02)** | | |
| Lys | VDW | K-Trp32, K-Gly91, K-Gln92, K-Ser93, K-Tyr94<br>H-Tyr100B |
| **Cys872 (22.88)** | | |
| Cys | VDW | H-Tyr100A, H-Tyr100B |
| Cys$^{N}$ | HB | H-Tyr100B$^{OH}$ |
| Cys$^{O}$ | HB | H-Tyr100B$^{OH}$ |
| **Ile873 (70.46)** | | |
| Ile | VDW | K-Trp32<br>H-Gly98, H-Tyr100A, H-Tyr100B |
| **Tyr874 (75.84)** | | |
| Tyr | VDW | H-Asn31, H-Tyr32, H-Ser96, H-Ser97, H-Gly98, H-Tyr100A |
| Tyr$^{O}$ | HB | H-Gly98$^{N}$ |
| Tyr$^{OH}$ | HB | H-Asn31$^{ND2}$ |
| **Arg875 (49.43)** | | |

| | | |
|---|---|---|
| Arg | VDW | H-Ser97, H-Gly98, H-Tyr99 |
| **Glu876 (33.92)** | | |
| Glu | VDW | H-Tyr32, H-Ser97 |
| $Glu^{OE1}$ | HB | H-$Tyr32^{OH}$, H-$Ser97^{OG}$ |
| **Tyr877 (16.19)** | | |
| Tyr | VDW | H-Tyr99 |
| $Tyr^{OH}$ | HB | H-$Tyr99^{OH}$ |
| **Asn885 (51.93)** | | |
| Asn | VDW | K-Asn28, K-Asn30, K-Gln92 |
| **Gln887 (17.10)** | | |
| Gln | VDW | K-Asn30 |

VDW: van der Waals
HB: hydrogen bond
SB: salt bridge

**Table S7. BSA (Å$^2$) by Ripr EGF**

| mAb | EGF6 | EGF6 linker | EGF7 | EGF7 linker | EGF8 | Total | EGF7/Total BSA (%) |
|---|---|---|---|---|---|---|---|
| **1G12** | 0 | 45.7 | 676.6 | 0 | 0 | 722.3 | 93.7 % |
| **2H1** | 372.2 | 59.9 | 200.4 | 0 | 0 | 632.6 | 31.7 % |
| **3F5** | 0 | 0 | 76.1 | 92.5 | 662.3 | 830.9 | 9.2 % |
| **5B3**[#] | 0 | 0 | 672.1 | 77.0 | 102.6 | 851.7 | 78.9 % |
| **7B3** | 86.0 | 22.6 | 590.9 | 0 | 0 | 699.5 | 84.5 % |

[#]Seager et al., 2026.

**Table S8. BSA (Å$^2$) by Fab CDRs**

| | H-chain | | | | L-chain | | | | |
|---|---|---|---|---|---|---|---|---|---|
| **mAb** | **CDR1** | **CDR2** | **CDR3** | **Total** | **CDR1** | **CDR2** | **CDR3** | **Total** | **Total** |
| **1G12** | 23.1 | 69.0 | 285.9 | 378.0 | 178.3 | 32.3 | 90.1 | 300.7 | 678.7 |
| **7B3** | 85.3 | 182.9 | 129.1 | 397.3 | 193.9 | 0 | 89.0 | 282.9 | 680.2 |
| **2H1** | 216.6 | 91.7 | 289.6 | 597.9 | 0 | 0 | 26.8 | 26.8 | 624.7 |
| **3F5** | 198.2 | 99.2 | 324.7 | 622.1 | 108.7 | 0 | 88.8 | 197.5 | 819.6 |

**Table S9. Cryo-EM data collection, refinement and validation statistics**

| **Data Collection** | **PfPCRCR–Fabs co-complex (EMDB-80770)** |
|---|---|
| Micrographs | 10,000 |
| Particles (Final map) | 68,128 |
| Pixel size (Å) | 0.639 |
| Defocus range (µm) | 0.5 – 1.6 |
| Voltage (kV) | 300 |
| Electron dose (e/Å$^2$) | 40 |
| Symmetry imposed | C1 |
| Initial particle images (no.) | 1,885,614 |
| Final particle images (no.) | 68,128 |
| Resolution (0.143 FSC) (Å) | 5.14 |
| Final Refinement | Non-Uniform Refinement |
| Map sharpening *B* | Not done |
| Model used | Crystal structures and AlphaFold3 predicted structures |
| Fitting | Rigid body |

**References**

1. Healer, J. *et al.* Neutralising antibodies block the function of Rh5/Ripr/CyRPA complex during invasion of *Plasmodium falciparum* into human erythrocytes. *Cell Microbiol.* **21**, e13030 (2019).
2. Seager, B.A. *et al.* PTRAMP, CSS and Ripr form a conserved complex required for merozoite invasion of Plasmodium species into erythrocytes. *Nat Commun* **17**, 1780 (2026).
